# Effective root responses to salinity stress include maintained cell expansion and carbon allocation

**DOI:** 10.1101/2022.09.01.506200

**Authors:** Hongfei Li, Kilian Duijts, Carlo Pasini, Joyce E van Santen, Nan Wang, Samuel C. Zeeman, Diana Santelia, Yanxia Zhang, Christa Testerink

**Affiliations:** Laboratory of Plant Physiology, Plant Sciences Group, Wageningen University&Research, 6708PB Wageningen, the Netherlands; Institute of Integrative Biology, ETH Zurich, 8092 Zurich, Switzerland; Institute of Molecular Plant Biology, ETH Zurich, 8092 Zurich, Switzerland

**Keywords:** carbon partitioning, cell expansion, halophytes, root growth, salt stress, *Schrenkiella parvula*

## Abstract

Acclimation of root growth is vital for plants to survive salt stress. Halophytes are great examples of plants that thrive under high salt concentrations but their salt tolerance mechanisms, especially those mediated by root responses, are still largely unknown. We compared root growth responses of the halophyte *Schrenkiella parvula* with its glycophytic relative species *Arabidopsis thaliana* under salt stress, and performed root transcriptomic analysis to identify differences in gene regulatory networks underlying their physiological responses. Primary root growth of *S. parvula* is less sensitive to salt compared with Arabidopsis. The root transcriptomic analysis of *S. parvula* revealed the induction of sugar transporters and genes regulating cell expansion and suberization under salt stress. ^14^C-labelled carbon partitioning analyses consistently showed that *S. parvula* had a higher incorporation rate of soluble sugars in roots under salt stress compared to Arabidopsis. Further physiological investigation revealed that *S. parvula* roots do not show a halotropic response and maintain root cell expansion and enhanced suberization even under severe salt stress. In summary, our study demonstrates that roots of *S. parvula* deploy multiple physiological and developmental adjustments under salt stress to maintain growth, providing new avenues to improve salt tolerance of plants using root-specific strategies.

## Introduction

Soil salinity is a serious environmental issue (Rengasamy, 2006). The loss of arable land and reduced crop production on salinity-affected soils challenge sustainable food production worldwide. While most crops are glycophytes, which are sensitive to salt, halophytes can survive and reproduce in environments with salt concentrations around 200mM NaCl or more (Flowers & Colmer, 2008; Orsini *et al.*, 2010; Kiani-Pouya *et al.*, 2017). Although salt response mechanisms of glycophytes have been extensively characterized, our understanding of the physiological processes that contribute to salt tolerance of halophytes is still in its infancy and potentially of great importance for engineering crop resilience.

Balancing sodium: potassium (Na^+^:K^+^) ratios and accumulation of compatible solutes or osmoprotectants are common strategies in plants to deal with salinity-induced ionic and osmotic stresses (Benito *et al.*, 2014; van Zelm *et al.*, 2020). Halophytes have developed unique strategies to cope with high salinity. For instance, the pseudocereal crop *Chenopodium quinoa* (*C. quinoa)* loads Na^+^ into its epidermal bladder cells through HIGH-AFFINITY POTASSIUM TRANSPORTER 1-type channels (Kiani-Pouya *et al.*, 2017; Bohm *et al.*, 2018). *Salicornia dolichostachya* and the Arabidopsis-relative halophyte *Eutrema salsugineum* (*E. salsugineum*) and *Schrenkiella parvula* (*S. parvula*) have constitutive high expression of gene *SALT OVERLY SENSITIVE 1* that encodes an ion transporter exporting Na^+^ from cells (Katschnig *et al.*, 2015; Tran *et al.*, 2021b). Genomic studies also found that both *C. quinoa* and *S. parvula* have a higher number of genes encoding sugar transporters compared to their glycophytic relatives (Dassanayake *et al.*, 2011; Zou *et al.*, 2017). *E. salsugineum* was reported to have constitutively higher levels of osmolytes such as soluble sugars, proline and gamma-aminobutyric acid compared to Arabidopsis, while *S. parvula* could rapidly increase and maintain abundant osmolytes and osmoprotectants under prolonged salt stress condition (Bartels & Dinakar, 2013; Tran *et al.*, 2021b). Although roots are the primary target of salt stress, the root adaptive strategies of halophytes are largely unknown.

Glycophytic plants have developed diverse adaptive root growth strategies to cope with salt stress. Upon salt perception, the rate of Arabidopsis root growth decreases and almost stops, followed by a partial recovery, resulting in slower growth rates overall, compared to control conditions (Geng *et al.*, 2013; Korver *et al.*, 2018). Furthermore, lateral root elongation is reduced more than that of the primary root (Julkowska *et al.*, 2014). *S. parvula* seedlings experienced much less inhibition of primary root growth by salt compared to Arabidopsis, and were also insensitive to treatment with the stress hormone, abscisic acid (ABA) (Tran *et al.*, 2021a; Sun *et al.*, 2022). At the cellular level, salt suppresses root growth by interfering with cell cycle progression in the root apical meristem (RAM) and by reducing elongation of mature cells (West *et al.*, 2004). Halotropism, facilitated by auxin redistribution, is a common strategy used by glycophytic species such as Arabidopsis and *Solanum lycopersicum*, by which the primary root actively grows away from high salt concentrations (Galvan-Ampudia *et al.*, 2013; Deolu-Ajayi *et al.*, 2019). The root endodermal layer serves as a checkpoint for radial transport of ions before they enter the stele. In Arabidopsis, the Casparian strip formed by lignin deposition encircles the endodermal cells and may play a role in limiting K^+^ leakage (Pfister *et al.*, 2014; Reyt *et al.*, 2021). Furthermore, suberin deposition between the endodermal plasma membrane and the cell wall is induced by salt to extend the coverage of endodermal cells limiting Na^+^ uptake from apoplast into the endodermis, hence contributing to the maintenance of Na^+^:K^+^ homeostasis (Barberon *et al.*, 2016). Despite the importance of adaptive strategies to sustain root growth under salinity, the contribution of root physiological and anatomical changes to plant salt tolerance of halophytes is still understudied.

In this study, we used *S. parvula* as a model halophyte species to investigate root developmental and growth response under salt stress in comparison with its glycophytic relative Arabidopsis. We combined transcriptomics and physiological analyses to decipher the molecular mechanisms underpinning root growth response of *S. parvula* under high salt conditions. We demonstrated that the relative insensitivity of *S. parvula* root growth under salt stress is reflected in a more efficient carbon partitioning to roots, maintenance of cell expansion and suberization, but not cell division. These strategies likely contribute to maintain root growth of *S. parvula* in saline environments.

## Materials and Methods

### Plant material and growth conditions

Seeds of *Arabidopsis thaliana* (Col-0) and *Schrenkiella parvula* (Lake Tuz ecotype) were stratified in dark at 4°C for 2 days and 7 days, respectively. For experiments performed on agar plates, seeds were surface-sterilized in 30% (v/v) sodium hypochlorite and washed 4 times with H2O. Seeds were geminated on ½-strength Murashige-Skoog (½MS) medium including vitamins (Duchefa), supplemented with 0.1% 2-(N-morpholino) ethanesulfonic acid (MES, Duchefa), 1% Daishin agar (Duchefa) (pH = 5.75). Unless otherwise specified, plates were placed at 70° racks in the growth chamber with 16h light:8h dark, 130μmol m^−2^ s^−1^ LED light, 21°C and 70% relative humidity.

### Analysis of the growth rate of primary roots

Four-day-old seedlings of *S. parvula* grown on vertical (90°) rack were transferred to new plates supplemented with 0mM, 125mM or 175mM NaCl and imaged every 2h for growth rate quantification (Gigli-Bisceglia *et al.*, 2022). The primary root length was quantified with Smartroot (Lobet *et al.*, 2011). Data points represent the average growth rate from 3 sequential timepoints. Nine rates were collected in 22h of monitoring. Calculation was performed with python-based script.

### Halotropism assay

*S. parvula* seeds were germinated on ½MS agar plates supplemented with 0.5% sucrose (basic medium) and transferred after 4 days to new plates where a 45° cut-out of the medium was made and replaced with basic medium supplemented with 0mM, 200mM, 300mM, 400mM or 500mM NaCl. Root tip positions were marked at 0h, 24h, 48h and 72h after transfer. Plates with 7-day-old seedlings were scanned with an Epson perfection V800 scanner at 300 dpi. Scanned images were transformed into greyscale and the primary root length and angle measured with Smartroot (Lobet *et al.*, 2011).

### Sample preparation and RNA sequencing

Root materials were harvested from *S. parvula* seedlings grown for 4 days on ½MS agar plates and transferred to new medium supplemented with 0mM or 175mM NaCl for 3h, 24h or 48h. Total RNA was isolated using NZY Total RNA Isolation kit (NZYtech). RNA library preparation and transcriptome sequencing were conducted by Novogene Co., LTD on a NovaSeq 6000 platform. Data will be deposited in the Array Express (http://www.ebi.ac.uk/arrayexpress) upon publication.

### Transcriptomic analysis

Read quality was assessed with FastQC (Andrews, 2010) and MultiQC (Ewels *et al.*, 2016) packages before and after trimming of low quality reads with Trim Galore! (Krueger, 2021). The reads were mapped to the transcriptome of *S. parvula* (Dassanayake *et al.*, 2011; Oh *et al.*, 2014) using Salmon (Patro *et al.*, 2017). Differential expression analysis was performed pairwise comparison of timepoints between treatments with DESeq2 (Love *et al.*, 2014). Differentially expressed genes (DEGs) were selected with the expression levels at |Log2 Fold Change|>1 and a false discovery rate (FDR) ≤0.01. For visualization in heatmaps the log fold changes were shrunk using “ashr” algorithm (Stephens, 2017). The heatmaps were generated with the R package pheatmap (Kolde, 2019). Comparative analysis on transcriptomes of roots of Arabidopsis (Geng *et al.*, 2013) and *S. parvula* was performed using k-means clusters with the Bioconda package “Clust” (Abu-Jamous & Kelly, 2018). FPKM values from the dataset of *S. parvula* and normalized microarray expression data of Arabidopsis were used (Geng *et al.*, 2013). Visualization was done with ggplot2 (Wickham, 2016). Gene Ontology enrichment was conducted using ClusterProfiler (Yu *et al.*, 2012; Wu *et al.*, 2021).

### ^14^CO_2_ Pulse-Chase Labeling

Ten-day-old seedlings of *S. parvula* and Arabidopsis were transferred to agar plates supplemented with 0mM or 150mM NaCl for 4h or 24h. The protocol of ^14^CO_2_ Pulse-Chase Labeling was adapted from (Kolling *et al.*, 2013; Thalmann *et al.*, 2016) and details were explained in Figure S1.

### Starch and soluble sugar measurements

Ten-day-old seedlings of *S. parvula* and Arabidopsis were transferred to agar plates supplemented with 0mM or 150mM NaCl for 4h or 24h. Shoots and roots were harvested and weighed. For shoot material: samples were homogenized in 0.7M perchloric acid and subject to centrifugation (10,000g for 5min at 4°C). The supernatant containing soluble sugars was adjusted to pH 6-7 with the buffer (2M KOH 400mM MES). For root material: samples were quenched in 0.8mL of 80% ethanol at 80°C for 10min then homogenized. After centrifugation (10,000g for 5min at 4°C), the pellet was sequentially washed with 0.5mL 50% ethanol, 20% ethanol and H2O. Supernatants were pooled, dried *in vacuo* and dissolved in 200ul H_2_O. Sugars (glucose, fructose, and sucrose) and starch were quantified enzymatically (Thalmann *et al.*, 2016).

### Cell wall and suberin staining

Seedlings were fixed in 4% paraformaldehyde in phosphate-buffered saline and cleared in ClearSee solutions (Ursache *et al.*, 2018). For cell wall staining, seedlings were stained in 0.1% Calcofluor White for 30min and washed in ClearSee for 30min. Suberin was stained with Nile Red (Sigma) and Fluorol Yellow 088 (FY 088, Sigma). For sequential staining of Nile Red and Calcofluor White: seedlings were stained with 0.1% Nile Red overnight, washed in ClearSee 3 times for 1 h, then stained with Calcofluor White as above. For FY 088 staining, seedlings were incubated in 0.01% FY 088 in methanol for 3 days, counterstained with 0.5% aniline blue (in H2O) for 1h, then rinsed in H_2_O (Berhin *et al.*, 2019).

### Microscopy

Microscopy experiments were performed either on a Leica TCS SP8 HyD confocal microscope (excitation and detection windows were as follows: Calcofluor White ex: 405nm, em: 425–475 nm; Nile Red ex: 561nm, em: 600–650nm) or a Leica DM5200 fluorescent microscope for roots stained with FY 088 (with a GFP filter ex: 450-490nm, em: 500-550nm). Images were analyzed with ImageJ. Four-day-old seedlings of Arabidopsis and *S. parvula* were transferred to agar plates supplemented with 0mM, 125mM or 175mM NaCl for 48h. Cortical cell length and cell number were quantified in roots of Calcofluor white-stained seedlings. Meristem size was measured as the distance from the quiescent center to the first cortical cell that is double the length of the previous cell. Cell length and number were quantified with ImageJ.

## Results

### Primary roots of *S. parvula* are insensitive to salt stress and contain an extra cortical cell layer

To investigate mechanisms of salt sensitivity during early seedling establishment, we examined root development and growth of *S. parvula* and Arabidopsis seedlings under salt stress. We transferred 4-day-old seedlings of *S. parvula* to salt gradients generated by media containing 200mM, 300mM, 400mM or 500mM NaCl at 1 cm from the root tip and measured the root growth direction and growth rate up to 72h after transferring. Interestingly, the root growth direction did not change compared to control conditions, independently of root growth rates (Fig. 1a-c, S2), suggesting *S. parvula*, unlike glycophytes, does not exhibit root halotropic responses upon salt perception. To understand the effect of salt on the root growth of *S. parvula*, we monitored primary root growth rate of *S. parvula* seedlings over 22h while exposed to 125mM or 175mM NaCl. The primary root growth was enhanced by 125mM NaCl after 10 h and was inhibited by 175mM NaCl (Fig. 1d). This contrasts to Arabidopsis primary root growth, which was strongly reduced under 125mM NaCl (Tran *et al.*, 2021a). Similar to salt stress, ABA treatments up to 15μM did not inhibit the primary root growth of *S. parvula* (Fig. S3), unlike Arabidopsis (Sun *et al.*, 2022).

**Figure 1.**
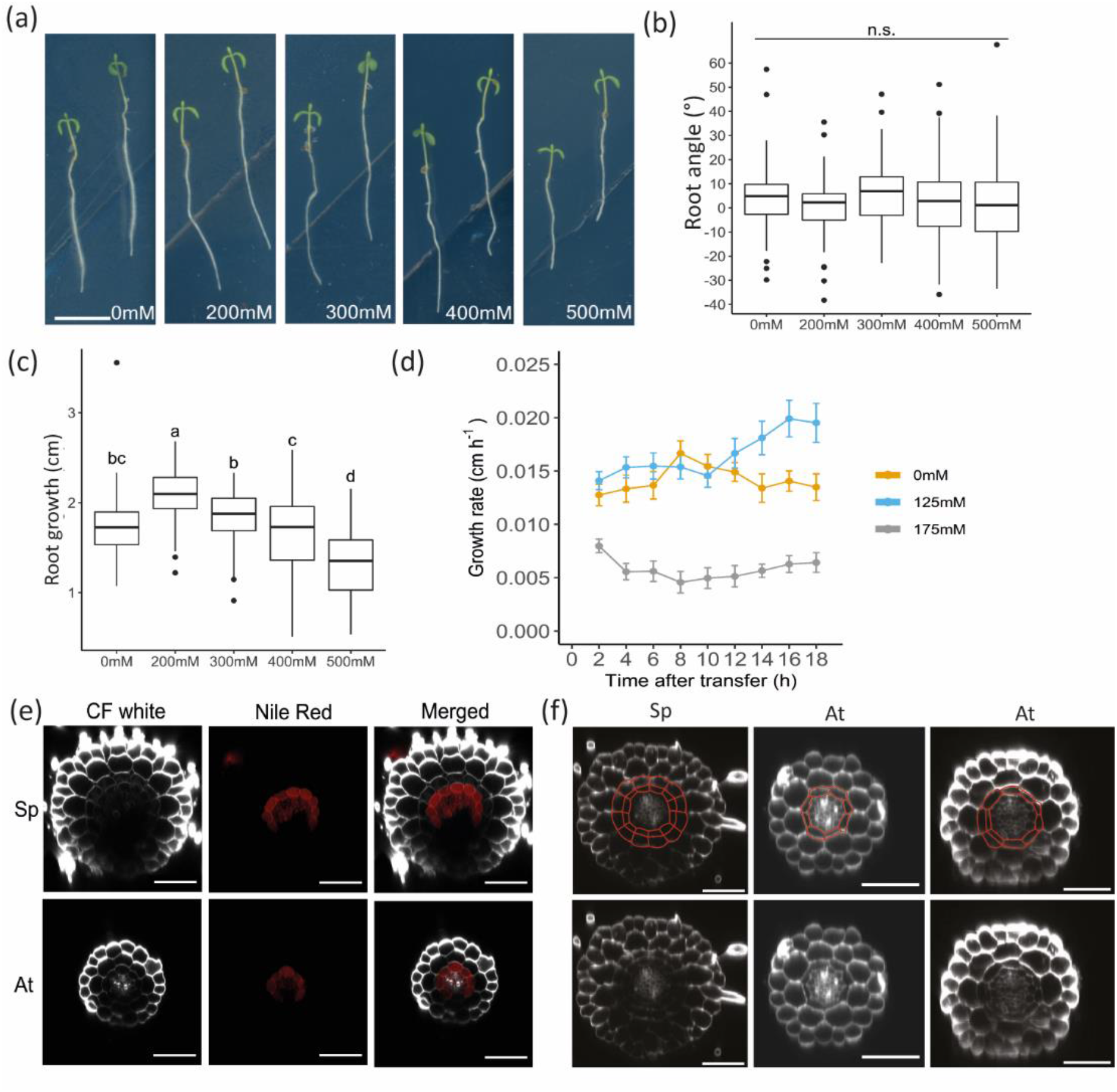
Root growth in response to salt and root patterning are different in *S. parvula* compared to Arabidopsis. (a, b, c) The roots of *S. parvula* do not actively avoid salt. Four-day-old *S. parvula* seedlings germinated and grown on ½ MS agar plates supplemented with 0.5% sucrose were transferred to replacement medium for 72h. Replacement medium (lower diagonal half) consisted of ½ MS medium supplemented with 0.5% sucrose and with 0, 200mM, 300mM, 400mM or 500mM NaCl. (a) Pictures of representative 4-day-old *S. parvula* exposed to diagonal salt gradients of 200-500mM NaCl for 72h. Scale bar=1cm. Primary root angle (b) and growth (c) of 4-day-old *S. parvula* seedlings exposed to diagonal salt gradient of 0, 200mM, 300mM, 400mM or 500mM NaCl for 24h (n=63-71). Significant differences (p<0.05) indicated by letters were determined by two-way ANOVA followed by Turkey’s post-hoc test and “n.s.” denotes no significant differences. (d) Growth rate of primary roots of *S. parvula* was monitored for 18h using a time-lapse setup. Four-day old *S. parvula* seedlings germinated and grown on ½ MS agar plates were transferred at time 0h to ½ MS medium supplemented with 0, 125mM or 175mM NaCl for 18h. Primary root length was quantified every 2h and each dot represents the average root elongation of every 3 time points. The individual data represents the average value of 26-30 seedlings from 3 independent experiments (n=26-30). Error bars represents the corresponding standard error of the mean. (e) The extra cell layer found in roots of 6-day-old *S. parvula* but not Arabidopsis, is not endodermis, but an extra cortex cell layer, as it lacks suberin. Four-day-old seedlings of *S. parvula* and Arabidopsis were transferred to ½ MS medium supplemented with 0, 125mM or 175mM NaCl for 48h. Cell walls were stained by Calcofluor White and suberin was stained by Nile Red. (f) Middle cortex is fully present in roots of 3-day-old seedlings *S. parvula* but not Arabidopsis. Cross sections were from the differentiation zone in roots of 3-day-old *S. parvula* and 3 and 7-day-old Arabidopsis. Seedlings of *S. parvula* and Arabidopsis were germinated and grown on ½ MS agar plates. Red lines in upper panel indicated the cell walls of endodermis and middle cortex and the original pictures were shown in lower panel. Scale bars=50μm.

To further investigate the morphological characteristics in roots that may contribute to salt tolerance, we examined root cross sections from *S. parvula* and Arabidopsis. We observed one extra layer in roots of 6-day-old *S. parvula* seedlings, but not in roots of Arabidopsis (Fig. 1e). To determine the identity of this extra cell layer, we used Nile Red to stain suberin, which usually locates in the endodermis of Arabidopsis roots (Barberon *et al.*, 2016). We observed that the extra layer was not stained (Fig. 1e), suggesting it is not likely the endodermis. It has been reported that a middle cortex (MC) initiates from endodermal cells of Arabidopsis roots 7 days after germination (Cui, 2016). We previously showed the extra layer in roots of *S. parvula* initiates from the endodermis (Tran *et al.*, 2021a), implying it could be MC. To further understand the initiation of this cell layer, we analyzed the presence of MC in different developmental stages of *S. parvula* and Arabidopsis seedlings. MC fully presented in roots of 3-day-old seedlings of *S. parvula* whereas it only partially formed in 6 to 7-day-old seedlings of Arabidopsis (Fig. 1f). We suggest an earlier appearance of MC in roots of *S. parvula* may be important for its halophytic lifestyle. To test whether MC formation is inducible by salt, we further investigated the MC presence in Arabidopsis under salt stress. However, MC presence was delayed rather than induced by salt (Fig. S4), which might be due to salt-induced global growth inhibition.

### Gene expression related to carbon partitioning in roots of *S. parvula* is affected by salt stress

To explore the molecular mechanisms underlying the salt response of *S. parvula* roots, we performed a transcriptomic analysis on roots of 4-day-old seedlings of *S. parvula* treated with or without 175mM NaCl for 3h, 24h or 48h. Principal component analysis (PCA) showed the relative relationships in the transcriptional profiles of salt-treated roots separated by treatments and timepoints. The transcriptional profile of root samples at 3h under salt stress was more separated from those of 24h and 48h (Fig. S5). The highest number of differentially-expressed genes (DEGs) was found at 3h, consisting of around 50% of DEGs of all time points (Fig. 2a, Table S1). These data suggest that salt stress induces a temporal reprogramming of gene expression in *S. parvula* roots, with a peak at 3h of stress.

**Figure 2.**
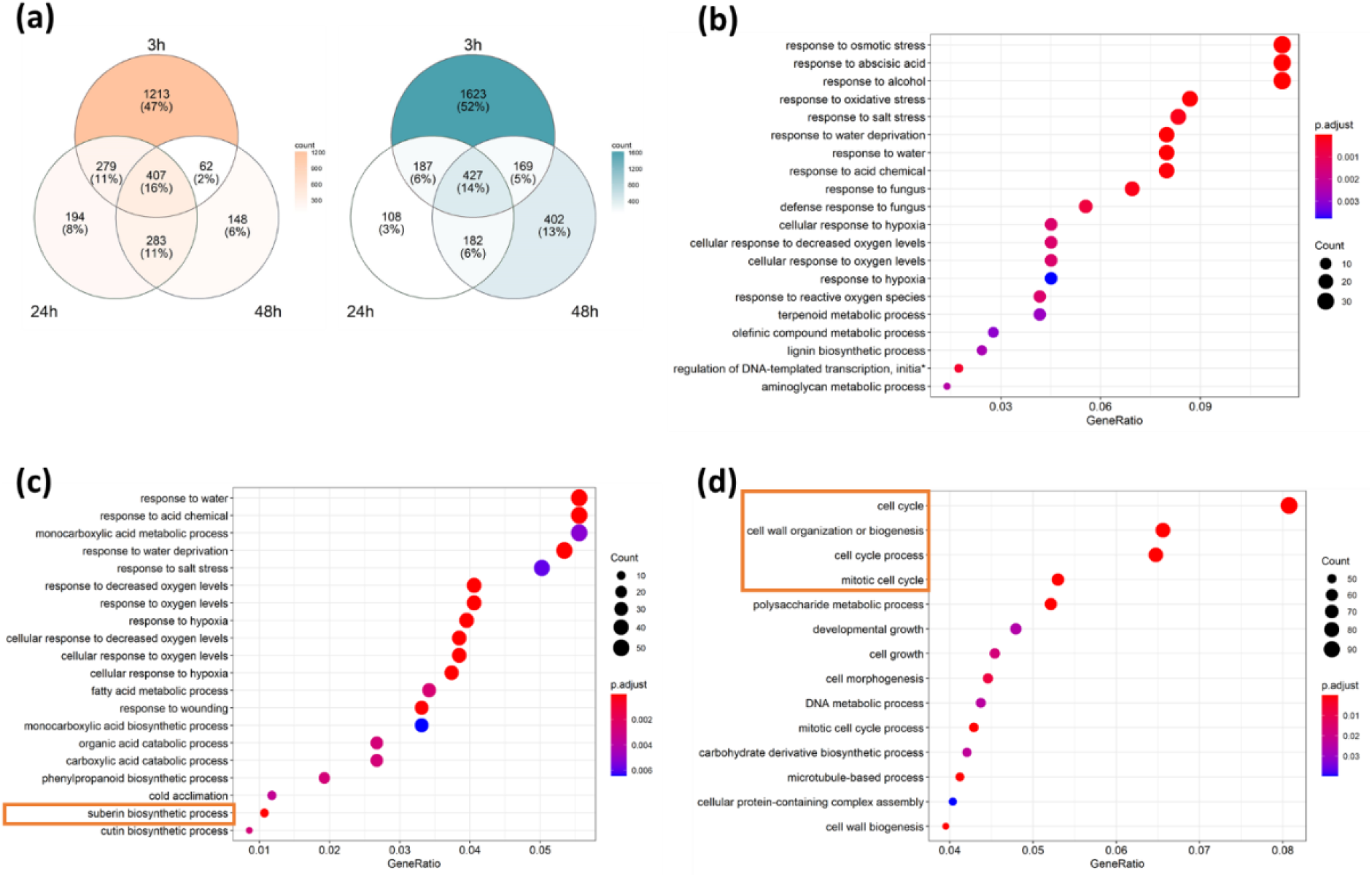
Temporal transcriptional responses of *S. parvula* roots under salt stress. Four-day-old *S. parvula* seedlings germinated and grown on ½ MS agar plates were transferred to ½ MS medium supplemented with 0 or 175mM NaCl for 3h, 24h and 48h. Roots were harvested for RNA sequencing. (a) Venn diagram presenting the number of significantly up- (pink) and down- (blue) regulated genes in roots of *S. parvula* at 3h, 24h and 48h after salt stress (|Log_2_ fold change| > 1; FDR < 0.01). (b) Gene ontology (GO) enrichment analysis of genes significantly up-regulated under salt stress at all timepoints (3h, 24h and 48h), based on biological process (BP). (c) GO enrichment analysis of genes only significantly up-regulated under salt stress at 3h, based on BP. (d) GO enrichment analysis of genes only significantly down-regulated under salt stress at 3h, based on BP.

Next, we performed gene ontology (GO) enrichment analysis on DEGs that are conserved in all timepoints and time-dependent. GO analysis on DEGs overlapping in all timepoints showed genes related to responses to diverse stimuli were constitutively upregulated from 3h to 48h (Fig. 2b, Table S2). Interestingly, similar terms were also enriched in GO analysis on DEGs that were only up-regulated at 3h but not at 24h and 48h (Fig. 2c and Table S4), indicating a wider-range of stress-responsive genes were induced at 3h. GO analysis on the DEGs that are only down-regulated at 3h included a number of biological processes related to cell cycle and growth, which may explain the inhibition of root growth under 175mM NaCl (Fig. 1d). Taken together, we found more transcriptional changes occurred at 3h, but responses to stress were activated in all timepoints.

Next, we investigated differences in key physiological traits between *S. parvula* and Arabidopsis, focusing on carbon partitioning, root suberization, and root cell elongation. First, we extracted expression patterns of *S. parvula* orthologs of Arabidopsis sugar transporters, including sucrose transporters (SUC) and SUGARS WILL EVENTUALLY BE EEXPORTED TRANSPORTED (SWEET) sucrose efflux transporters (Dassanayake *et al.*, 2011; Oh *et al.*, 2014). We also analyzed the well-established starvation marker *DARK-INDUCED6/ASPARAGINE SYNTHETASE1 (DIN6/ASN1)* (Baena-Gonzalez *et al.*, 2007; Muralidhara *et al.*, 2021). A strong induction of expression of *DIN6* was observed at 3h and the induction weakened at 24h, followed by a strong inhibition at 48h (Fig. 3a), suggesting a starvation response in roots of *S. parvula* under salt stress was induced at 3h but attenuated from 24h onwards. The expression of *SUC2* and *SWEET11* was strongly induced by salt stress at 3h and the expression of *SWEET11-15* was constantly elevated at 24h and 48h (Fig. 3a). AtSWEET11/12 and AtSUC2 mediate sugar loading in shoots in Arabidopsis and may also be responsible for apoplastic sugar unloading in roots given their high expression there (Gottwald *et al.*, 2000; Chen *et al.*, 2012; Durand *et al.*, 2016). Therefore, based on our gene expression data we hypothesize that roots of *S. parvula* may be rescued from salt-induced starvation after 24h due to enhanced sugar unloading in their roots.

**Figure 3.**
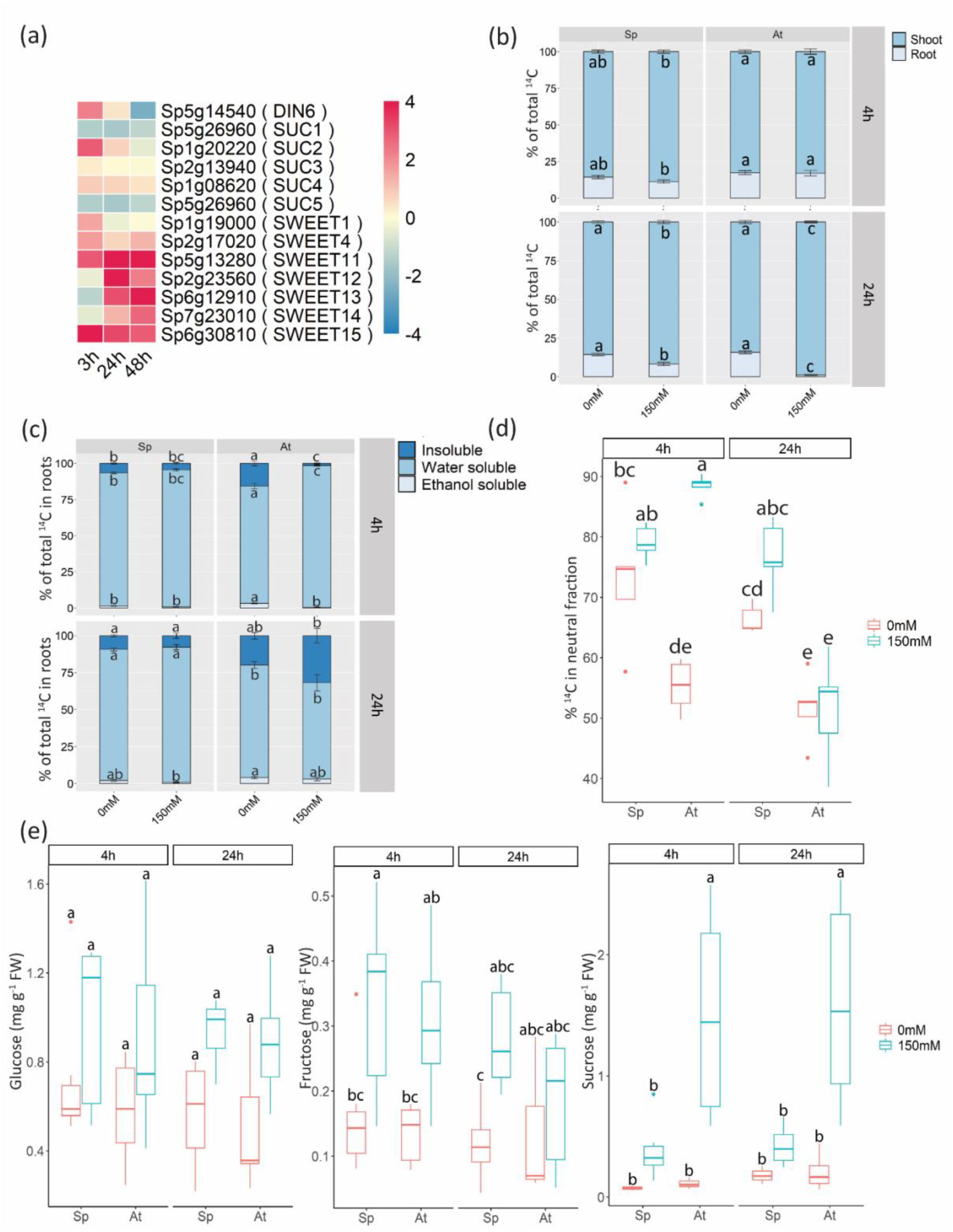
Better maintenance of carbon partitioning towards roots in *S. parvula* under salt stress. (a) Heatmap presenting expression profiles of starvation marker *DARK-INDUCED6* and detectable sugar transporters in roots of *S. parvula* under salt stress at 3h, 24h and 48h (columns). Gene expression visualization was extracted from transcriptomic data analysis. Gene symbols based on the annotations of Arabidopsis are shown in brackets (rows). Colors indicate the Log_2_ Fold changes between −4 and 4. (b) Carbon export to roots in *S. parvula* and Arabidopsis at 4h and 24h under salt stress. 10-day-old seedlings of *S. parvula* and Arabidopsis grown on ½ MS medium were transferred to ½ MS medium supplemented with 0mM or 150mM NaCl for 4h or 24h. Incorporation of ^14^C into shoots and roots was presented as percentages of total ^14^C in plants (n= 5-6 pools of 10-15 roots or shoots). (c) The incorporation of ^14^C into different compounds in roots at 4h and 24h under salt stress. The amount of incorporated ^14^C into all compounds was presented as percentages of total ^14^C in roots (n= 5-6 pools of 10-15 shoots or roots). (d) Carbon incorporation into neutral fraction *i.e*. soluble sugars from water-soluble compounds under salt stress in roots at 4h and 24h under salt stress. The amount of incorporated ^14^C into the soluble sugars in roots was given as percentages of incorporated ^14^C in water-soluble compounds. (e) Soluble sugars (glucose, fructose and sucrose) content in roots of *S. parvula* under salt stress. 10-day-old seedlings of *S. parvula* and Arabidopsis grown on ½ MS medium were transferred to ½ MS medium supplemented with 0mM or 150mM NaCl for 4h or 24h. FW, fresh weight. Significant differences (p<0.05) indicated by letters were determined by two-way ANOVA followed by Turkey’s post-hoc test.

To test this hypothesis, we conducted ^14^CO_2_ pulse-chase labeling experiments to trace carbon partitioning from shoots to roots and into different compounds under salt stress. Under control conditions, carbon partitioning in *S. parvula* and Arabidopsis was similar, with over 80% of total amount of ^14^C fixed by plants found in the shoots (Fig. 3b). Interestingly, carbon export to roots was slightly reduced after 4h salt treatment in *S. parvula* but not in Arabidopsis (Fig. 3b), which may indicate an earlier response to salt in *S. parvula*. After 24h exposure to salt, carbon allocation to roots was reduced to almost zero in Arabidopsis, whereas export was maintained in *S. parvula*, albeit at a lower rate than control conditions. This suggests that *S. parvula* has a stress-response mechanism enabling it to continue allocating carbon to its roots under salt stress that Arabidopsis lacks. Subfractionation of the fixed ^14^C in roots and shoots revealed differences in the carbon partitioning between *S. parvula* and Arabidopsis. The allocation of ^14^C incorporation into water-soluble compounds (*i.e*., amino acids, organic acids, sugar phosphates and neutral sugars) was higher in roots and shoots of *S. parvula* under both control and salt conditions and carbon partitioning dynamics were only mildly disturbed by salt (Fig. 3c,d and S6). In contrast to *S. parvula*, Arabidopsis increased ^14^C incorporation into water-soluble compounds in shoots in response to salt at 4h and 24h, while in roots only at 4h (Fig. 3c and S6). As neutral soluble sugars are an important source of energy and osmolytes, we further isolated neutral compounds from the water-soluble fraction. In *S. parvula* roots, over 60% of water-soluble compounds were in the neutral fraction (*i.e*. soluble sugars). This proportion increased slightly upon salt treatment (Fig. 3d). In Arabidopsis roots, 50% of water-soluble compounds were in the neutral fraction under control conditions. After 4h salt treatment, this fraction had increased close to 100%, suggesting that carbon imported into the root was not metabolized normally. After 24h treatment, when the export rate in Arabidopsis was greatly reduced, the proportion of ^14^C in the neutral compounds had returned to control levels (Fig. 3d). These results suggest not only that partitioning of carbon to roots upon salt stress is more stable in *S. parvula*, but also that its metabolism is less disrupted than in Arabidopsis. Next, we measured the soluble sugar pools (glucose, fructose and sucrose) in roots of *S. parvula* and Arabidopsis using enzymatic assays. The response to salt treatment was modest in *S. parvula* with a tendency towards slightly increased levels of all three sugars (Fig. 3e). In Arabidopsis, however, sucrose levels increased substantially in roots and was manifold higher than in control values after 24h salt treatment (Fig. 3e). These are in line with less perturbation of carbon partitioning in roots of *S. parvula* under salt stress. In shoots of *S. parvula*, the fraction of neutral compounds from the water-soluble compounds stayed stable after 4h and 24h salt treatments while the incorporation into the water-soluble fraction slightly increased after 24h salt treatment (Fig. S6). In shoots of Arabidopsis, the fraction of salt induced neutral compounds from the water-soluble fraction increased to over 60% after 4h and dropped after 24h while the total incorporation into the water-soluble fraction was elevated to over 90%. The measurements on sugar pools revealed both species accumulated more soluble sugars under stress conditions (Fig. S6c). These data are in line with the carbon partitioning in roots that most assimilated carbon was incorporation into water-soluble, mainly neutral sugars in Arabidopsis in response to salt stress while it is less interrupted in *S. parvula*.

### *S. parvula* can maintain suberization under severe salt stress

Suberization in the root endodermis is considered a barrier to control radial ion transport to the stele (Barberon *et al.*, 2016). Measurements of Na^+^ and K^+^ in 4-day-old seedlings after 2-day exposure to 0, 125mM or 175mM NaCl showed that *S. parvula* accumulated less Na^+^ than Arabidopsis (Fig. S7a). Although the overall K^+^ content was reduced by salt, *S. parvula* had higher levels of K^+^ under control conditions and less reduction under 125mM and 175mM NaCl, resulting in much lower Na^+^:K^+^ ratios in *S. parvula* compared to Arabidopsis (Fig. S7b, c). These data suggest that *S. parvula* seedlings maintain a favorable ion balance by limiting Na^+^ influx and maintaining high K^+^ levels upon salt treatment, as was found for adult *S. parvula* plants (Orsini *et al.*, 2010; Tran *et al.*, 2021b). Interestingly, we found the GO term ‘suberin biosynthetic process’ was enriched by the up-regulated DEGs at 3h (Fig. 2c). The expression pattern of genes regulating suberin biosynthesis reveal that transcription factors of MYeloBlastosis (MYB) family (*MYB39, 42, 53, 92 and 93*), suberin biosynthesis and polymerization genes were induced in *S. parvula* at 3h under salt stress (Fig. 4a). Thus, we measured the root suberization of 4-day-old seedlings of *S. parvula* and Arabidopsis after 6h and 24h exposure to 125mM or 175mM NaCl. In both species, the suberization zone starts from the differentiation zone and ends close to the root tip, which includes a fully-formed suberization zone and discontinuous suberization (patchy) zone (Fig. 4b, c). Under control conditions, *S. parvula* had a shorter fully-suberized zone than Arabidopsis (Fig. 4c, S8) and, at 6h salt treatments (125mM or 175mM NaCl), suberin coverage in both species was unchanged (Fig. S8). After 24h salt treatments, suberin coverage was greatly increased in *S. parvula* due to an increased full-suberized zone (Fig. 4c, 4d). In Arabidopsis, the situation differed: suberin coverage was increased under 125mM NaCl but was reduced under 175mM NaCl compared to control condition (Fig. 4c, 4d). This was consistent with the lethality of 175mM NaCl to Arabidopsis (Tran *et al.*, 2021a). We further compared relative changes of suberization under salt stress and found the fully-suberized zone in *S. parvula* increased around 3 times in length under 125mM NaCl, and even further under 175mM NaCl, whereas it increased less than 2 times under 125mM NaCl and did not increase under 175mM NaCl in Arabidopsis (Fig. 4d). These results revealed that *S. parvula* is capable to increase suberization under severe salt stress (175mM NaCl) in roots, which may also contribute to the better ion balance (Fig. S7).

**Figure 4.**
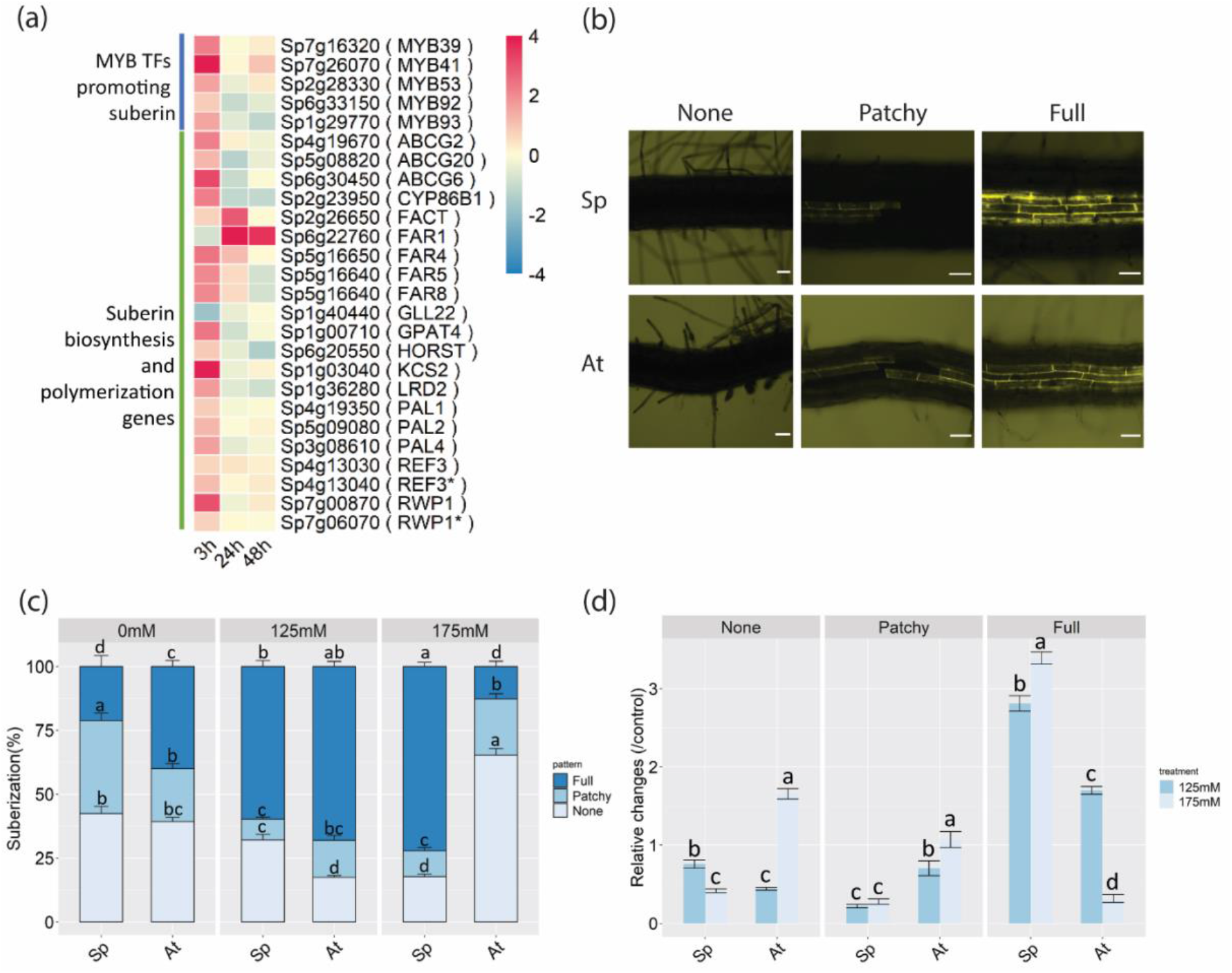
More suberization in response to salt in roots of *S. parvula* compared to Arabidopsis. (a) Heatmap presenting expression profiles of genes involved in suberin biosynthesis (Shukla *et al.*, 2021) in 3h, 24h and 48h (columns) under salt stress in roots of *S. parvula*. Gene expression was extracted from transcriptomic data analysis. Gene symbols based on the annotations of Arabidopsis were shown in brackets (rows). Colors indicate the Log_2_ Fold changes between −4 and 4. (b) Images of three different suberization zones of *S. parvula* (Sp) and Arabidopsis (At) roots. Analysis of suberization in roots of *S. parvula* under salt stress was performed on 4-day-old seedlings of *S. parvula* and Arabidopsis were transferred to ½ MS medium supplemented with 0.5% sucrose and with 0, 125mM or 175mM NaCl for 24h and suberin was stained by FY088. Suberization quantification was separated into non-suberized zone (None, left), patchy (Patchy, middle) and continuous suberization (Full, right). Scale bars=50μm. (c) Quantification of suberin deposition along the root axis and was indicated with 3 different zones (n=23-27). Data are presented as the percentage of root length. Error bars represent standard error. Significant differences (p<0.05) indicated by letters were determined by two-way ANOVA followed by Turkey’s post-hoc test. (d) Quantification of relative changes of 3 suberization zones under salt stress. Data are calculated by the ratio between salt treatment and control condition. Error bars represent standard error. Significant differences (p<0.05) indicated by letters were determined by two-way ANOVA followed by Turkey’s post-hoc test.

### Fast recovery of cell expansion ensures root growth under salt stress

Primary root growth of *S. parvula* was less inhibited under 175mM NaCl which is lethal to Arabidopsis (Fig. 1d) (Tran *et al.*, 2021a; Gigli-Bisceglia *et al.*, 2022). However, GO analysis revealed that genes involved in processes underpinning root growth, i.e. ‘cell cycle’ and ‘cell growth’, were inhibited in roots of *S. parvula* 3h after application of 175mM NaCl (Fig. 2d). Therefore, we further analyzed the expression of these DEGs at subsequent timepoints. Interestingly, DEGs that were down-regulated at 3h and annotated with the term ‘cell cycle’ were not significantly down-regulated after 24h or 48h (Table S3), suggesting salt stress imposed only transient inhibition of the cell cycle in *S. parvula* roots. Moreover, a subset of genes from the down-regulated DEGs enriched in GO term ‘cell growth’ were upregulated more than 2 times from 24h or/and 48h, including *UNS2* and several expansins (*EXPANSIN A4* (*EXPA4*), *EXPA5*, *EXPA6, EXPA12*) (Fig. 5a). *UNS2*, also known as *HOOKLESS1*, is required for root elongation (Lehman *et al.*, 1996) and expansins are cell wall loosening enzymes which are necessary for cell expansion (Feng *et al.*, 2016; Majda & Robert, 2018). The GO term ‘cell wall organization or biogenesis’ was enriched in upregulated DEGs at 24h and 48h (Fig. S9, Table S3), which includes cell wall biosynthetic enzymes that supply cell wall components for long-term cell growth (Cosgrove, 2014; Cosgrove, 2016; Feng *et al.*, 2016) (Fig. S9). These data suggest that the cell growth in roots of *S. parvula* is not strongly inhibited by salt treatment due, in part, to transcriptomic reprogramming.

**Figure 5.**
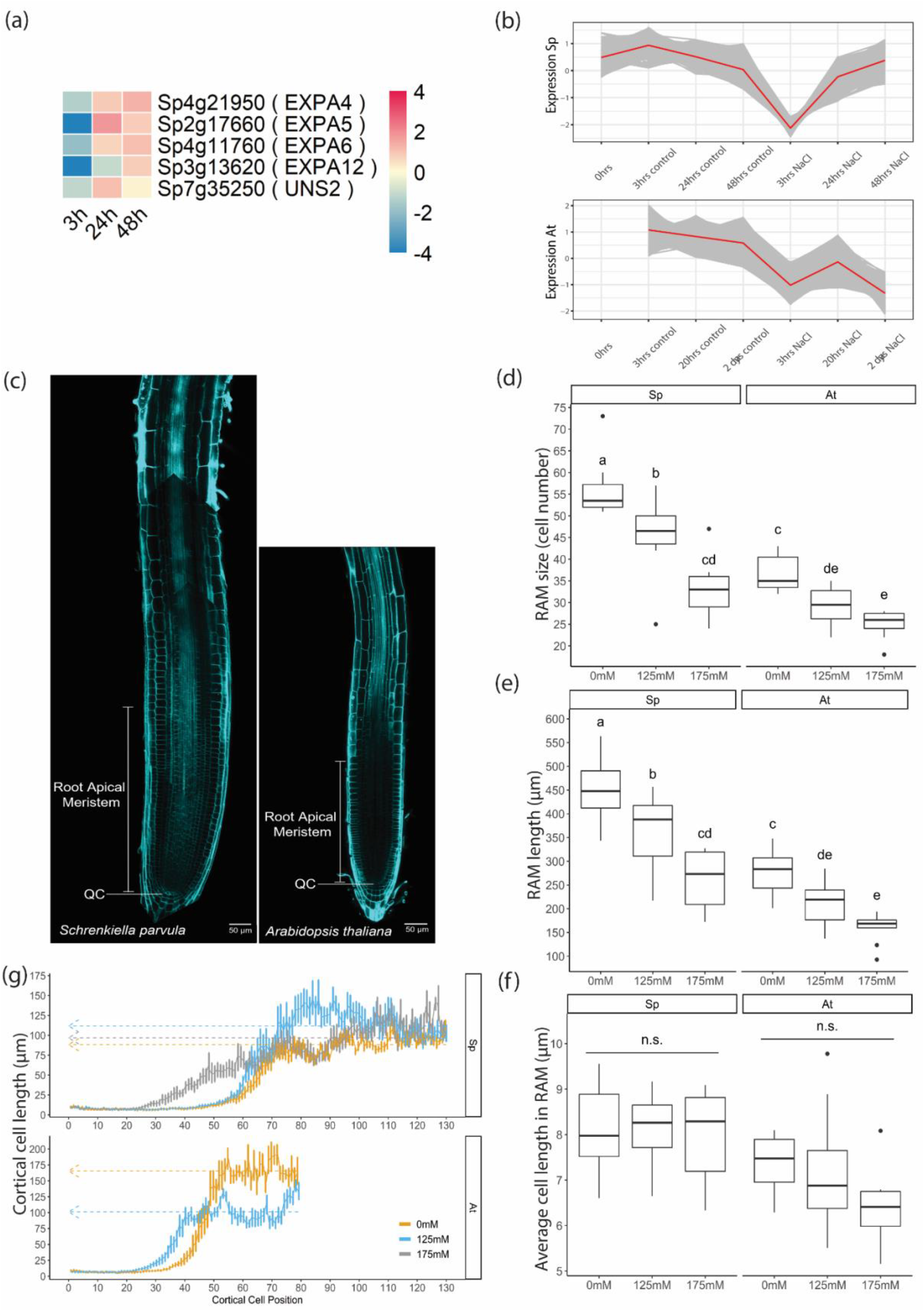
Exposure to salt does not inhibit cell expansion but reduces cell division in roots of *S. parvula*. (a) Heatmap presenting expression profiles of DEGs down-regulated in GO term ‘cell growth’ at 3h but up-regulated (*p*< 0.05 Log_2_ Fold changes >1) at 24h and/or 48h in roots of *S. parvula* under salt stress. Gene expression was extracted from transcriptomic data analysis. Gene symbols based on the annotations of Arabidopsis were shown in brackets (rows). Colors indicate the Log_2_ Fold changes between −4 and 4. (b) Expression profiles of genes in Cluster 1 from K-means cluster analysis. K-means cluster analysis was performed on transcriptomes of roots of *S. parvula* (upper panel) and public data from Arabidopsis roots (Geng *et al.*, 2013) (lower panel). Red lines indicate the average of the expression. (c) Illustration of the quantification of root meristem size. Primary roots of 4-day-old *S. parvula* and Arabidopsis transferred to new plates of ½ MS medium for 2 days. QC represents the quiescent center where the quantification of root meristem starts. White arrows indicate root meristem size in roots. Scale bars = 50 μm. (d-f) Root apical meristem (RAM) sizes and cell numbers but not cell length of *S. parvula* and Arabidopsis are decreased by salt. RAM length (d), number of cells (e) and average cell length in RAM (f) were quantified (n=8-15). Four-day-old seedlings of Arabidopsis and *S. parvula* germinated and grown on ½ MS agar plates were transferred to ½ MS agar plates supplemented with 0mM, 125mM or 175mM NaCl for 48h. Error bars represent standard error. Significant differences (p<0.05) indicated by letters were determined by two-way ANOVA followed by Turkey’s post-hoc test. (g) Cell expansion is inhibited in Arabidopsis roots by salt, but not in roots of *S. parvula*. Cortical cell length was measured from quiescent center (position 0) towards shoots (n=8-10). Striped arrow lines denote average cortical cell length in differentiation zone.

We further compared transcriptomics data on the root salt response of *S. parvula* with public dataset of transcriptomes from the roots of Arabidopsis under salt stress (Geng *et al.*, 2013) using K- means clustering (Abu-Jamous & Kelly, 2018). The expression profiles were clustered into 9 sets (Fig. 5b, S10). Clusters 0, 5, 7 and 8 showed similar expression patterns between *S. parvula* and Arabidopsis, suggesting that there are common underlying stress responses between *S. parvula* and Arabidopsis. Several distinct transcriptional changes were also observed. Genes in cluster 1 had similar expression profiles in *S. parvula* and Arabidopsis at 3h, with a general decrease in transcript abundance; but their expression diverged at the 2-day/48h timepoint (Fig. 5b). Interestingly, the GO analysis of biological processes performed on the genes in Cluster 1 showed an enrichment of terms related to cell division and expansion such as ‘cell cycle’ and ‘microtubule cytoskeleton organization’ (Fig. S11 and Table S6) (Li *et al.*, 2015; Qi & Zhang, 2020). This may indicate a faster recovery of *S. parvula* roots than Arabidopsis from the inhibition of cell division and expansion caused by salt.

To further assess how salt affects root elongation at cellular level, we compared cell division and cell expansion in the primary roots of *S. parvula* and Arabidopsis under salt stress. We quantified RAM size by measuring both cell number and RAM length, and found that both *S. parvula* and Arabidopsis had a reduced RAM size in response to salt, even though root growth of *S. parvula* was promoted under 125mM NaCl (Fig. 1d, 5c-e). The average cell length of the RAM indicated that cell number rather than cell length was inhibited under 125mM and 175mM NaCl in both species (Fig. 5f). Moreover, to understand how cell expansion in primary roots of *S. parvula* is affected by salt, we measured cell length from the quiescent center (QC) to the differentiation zone. Consistent with the observation that cell length in RAM is not inhibited by salt in either species, the cortical cell length stayed low and stable from QC in both species independent of salt concentrations (Fig. 5g). Under 125mM NaCl, cortical cell length of both *S. parvula* and Arabidopsis roots started increasing at the position closer to root tips which is in line with their smaller RAM sizes in terms of cell number under 125mM NaCl (Fig. 5d, 5e). However, cell length of *S. parvula* kept increasing under 125mM NaCl until it was almost 1.5 times longer than cells at same position under control condition, whereas cell length in Arabidopsis was strongly reduced (Fig. 5g). Under 175mM NaCl, which is lethal to Arabidopsis seedlings in our experiment, cell length in primary roots of *S. parvula* showed similar changes as control condition after 65 cells. In summary, the cellular mechanism that allows the roots of *S. parvula* to keep growing under salt stress appears due to its capability to maintain cell expansion, but not cell division.

## Discussion

The plasticity of root growth supports plant survival in saline conditions. Halophytes are great models to understand how plants can thrive under high salinity, potentially unravelling novel salt stress tolerance mechanisms. Hereby, we reported that, unlike Arabidopsis, roots of *S. parvula* did not grow away from higher salt concentrations and their root growth was less inhibited by salt and ABA. Based on a transcriptomic survey on roots of *S. parvula* under salt stress and further physiological analyses, we found that several characteristics of *S. parvula* may contribute to its root growth in saline enviroments, including strategies of carbon partitioning, maintenance of cell expansion and suberization.

### Do root anatomical characteristics contribute to stress resilience?

We observed that the appearance of MC was earlier in roots of *S. parvula* than in Arabidopsis (Fig. 1e). Notably, *S. parvula* is not the only Arabidopsis relative having different root patterning. Roots of *E. salsugineum* have one extra layer of both cortex and endodermis at adult-stage (Inan *et al.*, 2004); Primary roots of *Cardamine hirsuta* have a pre-patterned middle cortex during embryogenesis (Di Ruocco *et al.*, 2018). Moreover, multiple cortical layers exist in many monocot plants such as barley, maize and rice (Alarcon *et al.*, 2014; Henry *et al.*, 2015; Kirschner *et al.*, 2017). In the roots of rice and barley, inner cortex layers next to endodermis have a flattened shape similar to endodermis and thicker cell walls, whereas outer cortex layers exhibit a rounder shape and have more air-containing cavities between cells (Henry *et al.*, 2015), which morphologically resembles the middle cortex and cortex in roots of *S. parvula* (Fig. 1e, 1f). The regulation of HOMEODOMAIN LEUCINE ZIPPER III (HD-ZIPIII) transcription factor *PHABULOSA* by microRNA 165 and 166 (miR165/6) in middle cortex/endodermis initial cells enables the additional periclinal division in *Cardamine hirsuta* leading to the middle cortex formation during late embryogenesis (Di Ruocco *et al.*, 2018). We found salt stress delayed middle cortex appearance in roots of Arabidopsis, rather than inducing it (Fig. S4), which may be due to the mis-regulation of miR165- HD-ZIPIII cascade by salt-induced changes in phytohormone signals including ABA and auxin (Li *et al.*, 2021). Even though multiple cortical layers are common to the roots of many species, our knowledge on the regulation and the function of multiple cortical layers under abiotic stress is still very limited. Single cell RNA sequencing techniques might be a great solution to unravel this.

*S. parvula* seedlings accumulated less Na^+^ and maintained higher K^+^ than Arabidopsis under salt stress (Fig. S7), suggesting the existence of an effective mechanism to limit Na^+^ influx and/or K^+^ leakage. Indeed, we found up-regulated DEGs related to suberin and lignin biosynthesis at 3h and a consistently increased suberin coverage in roots of *S. parvula* under severe salt stress at 3h (Fig. 4c, 4d and Table S4). Suberization and lignification in endodermal cells of roots function as barriers to control the radial transport of ions into the stele (Barberon *et al.*, 2016; Reyt *et al.*, 2021). Suberin is induced by ABA and salt stress to coat the endodermal cells in roots and thus limits the coupled transcellular pathways, whereas Casparian strip mainly made of lignin controls apoplastic pathways. Defects in suberization and/or lignification trigger the over-accumulation of Na^+^ and reduction of K^+^ (Barberon *et al.*, 2016; Reyt *et al.*, 2021). This indicates that the maintenance of suberization in the roots of *S. parvula* may contribute to its ion homeostasis regulation and possibly leads to less growth reduction in response to salt stress.

### Carbon partitioning in roots of *S. parvula* is less affected by salt stress than in Arabidopsis

Limited carbon fixation under abiotic stress due to a reduced photosynthesis rate, decreased stomatal conductance and restricted CO_2_ uptake may challenge carbon export to roots (Munns, 2002). Reduced CO_2_ assimilation and stomatal conductance were observed under 50mM NaCl in 8-week-old Arabidopsis plants but only under 150mM NaCl in 8-week-old *S. parvula* although the basal levels of CO_2_ assimilation and stomatal conductance were lower in *S. parvula* than in Arabidopsis (Tran *et al.*, 2021a). Consistently, carbon export to roots and metabolism in roots of *S. parvula* experienced less disturbance by salt compared with Arabidopsis (Fig. 3). Consistently, 5-week-old Arabidopsis plants were also shown to allocate less carbon towards the roots after 24h of salt treatments (100mM and 200 mM NaCl) (Dong *et al.*, 2018). However, water deficit was shown to enhance carbon export to the roots of Arabidopsis (Durand *et al.*, 2016). The difference in carbon partitioning between salt stress and water deficit may be due to salt-induced ion toxicity (van Zelm *et al.*, 2020). In our study, the higher ability of *S. parvula* in maintaining carbon homeostasis in the roots under salt stress, is likely to contribute to the less inhibited root growth of *S. parvula* than Arabidopsis under salt stress (Fig. 1c, 1d).

In *S. parvula* roots, the carbon incorporation into different compounds and soluble sugars in water-soluble compounds after 4h and 24h salt treatments were maintained at comparable levels as control conditions (Fig. 3c, 3d). In Arabidopsis roots, almost all carbon stayed in water-soluble compounds after 4h and was back to comparable levels as control after 24h (Fig. 3c, 3d). Carbon fixation via photosynthesis produces sugars to transport to roots. Therefore the predominantly high fraction in soluble sugars in roots of Arabidopsis under salt stress suggests they are maintained in the form in which they arrive roots. Consistently we observed higher levels of sucrose in roots of Arabidopsis after 4h and 24h salt treatments (Fig. 3e). These may be actively regulated, to serve as an internal osmolyte protecting cells from osmotic stress imposed by salinity (Slama *et al.*, 2015). As *S. parvula* accumulates less Na^+^ (Fig. S7), it may experience less stress than Arabidopsis and sucrose demand as osmolytes may be lower. Alternatively, it could be a passive response due to a lack of conversion of imported sugars into other forms. If central metabolism is inhibited by salt, sugars will not be converted into other respiratory intermediates, and starvation will ensue. Indeed, Na^+^ interferes with diverse enzymatic processes that require K^+^ due to their chemical similarity (van Zelm *et al.*, 2020). The over-accumulation of salt may inhibit sucrose metabolism more in Arabidopsis than in *S. parvula* due to its better ion balance (Fig. S7). Moreover, a constant up-regulation of the starvation marker *ASN1* and increasing levels of sucrose were observed in roots of Arabidopsis after 6h salt treatment (Hartmann *et al.*, 2015), suggesting an imbalance of sucrose metabolism and energy supply in salt-treated roots of Arabidopsis. Taken together, *S. parvula* is likely to have a more stable carbon partitioning and sugar metabolism under salt stress helping its survival under salt stress.

### Distinct growth responses in roots of *S. parvula* may be due to differences in ABA- and auxin- related hormonal signaling networks

Primary root growth of *S. parvula* was less sensitive to salt stress and ABA treatments compared with Arabidopsis (Fig. 1d, S2), which is consistent with a recent report (Sun *et al.*, 2022). ABA biosynthesis and signaling pathways are activated by diverse abiotic stresses including salinity (Vishwakarma *et al.*, 2017). ABA represses root growth through interactions with other phytohormones including auxin and ethylene. ABA-triggered ethylene biosynthesis and signaling induces auxin biosynthesis in root meristems and transport to elongation zones to inhibit cell elongation (Stepanova *et al.*, 2005; Ruzicka *et al.*, 2007; Swarup *et al.*, 2007; Stepanova *et al.*, 2008; Strader *et al.*, 2010; Luo *et al.*, 2014). DNA affinity purification (DAP-Seq) and transcriptomic analysis revealed that *S. parvula* lost the regulation of *ABA-RESPONSIVE ELEMENT BINDING FACTORS* (*AREB/ABFs*) of the auxin signaling gene *INDOLEACETIC ACID-INDUCED PROTEIN 16* (*IAA16*) compared to Arabidopsis, which expression was not changed in roots of *S. parvula* in response to ABA (Sun *et al.*, 2022). Moreover, AREB/ABFs-targeted ethylene biosynthesis enzyme *ACC synthase 2* and *7* and several auxin-related genes such as auxin efflux carrier *PIN1, PIN7* and *IAA11* showed different transcriptional regulations by ABA in *S. parvula* compared with Arabidopsis (Sun *et al.*, 2022). This may imply that *S. parvula* has different regulation of auxin pathways in response to ABA, leading to the observed insensitive root growth of *S. parvula* under ABA and salt treatments. Consistently, we showed that roots of *S. parvula* failed to exhibit a halotropic response even when root growth was significantly inhibited by salt (Fig. 1a-c). Halotropism is conducted through asymmetric auxin distribution via endocytosis of PIN2 (Galvan-Ampudia *et al.*, 2013). ABA reduces PIN2 abundance mainly through the inhibition of *PIN2* expression in roots of Arabidopsis (Xie *et al.*, 2021). Interestingly, the expression of *PIN2* was more inhibited in roots of *S. parvula* than Arabidopsis under ABA treatments (Sun *et al.*, 2022). Thus, a different regulation of *PIN2* by ABA may play a role in the non-halotropic response in roots of *S. parvula*.

### Roots of *S. parvula* maintained cell expansion possibly due to changes in cell wall loosening and biosynthesis

Cell wall plasticity helps protect cells from external stimuli but also accommodating the increase in cell size (Vaahtera *et al.*, 2019). The increased cell wall extensibility by cell wall loosening proteins and enzymes such as expansins, xyloglucan endotransglycosylase/hydrolase (XTH) and cell wall biosynthesis to counteract cell wall breakage caused by overextension, are required for cell expansion (Cosgrove, 2000; Van Sandt *et al.*, 2007; Cosgrove, 2016). Our transcriptomic data indicated that the upregulation of genes encoding cell wall loosening proteins/enzymes and putative non-cellulosic polysaccharides synthesis enzymes from 24h onwards may contribute to the maintenance of cell expansion in roots of *S. parvula* under salt stress (Fig. 5a, S9, Table S3, S4). Moreover, the GO analysis on DEGs upregulated at 24h and 48h in roots of *S. parvula* were annotated with the GO term ‘cell wall organization or biogenesis’ (Table S6). However, in Arabidopsis, genes related to structural constituents of the cell wall were down-regulated in most root layers except stele at 20h and 2 days under salt stress (Geng *et al.*, 2013). These may indicate an active cell wall biosynthesis in roots of *S. parvula* in prolonged salt stress, but not in Arabidopsis. Consistently, DAPseq and RNAseq analysis also showed that several genes encoding cell wall loosening- and biosynthesis-related enzymes regulated by ethylene or auxin were differentially regulated by ABA in roots of *S. parvula* (Sun *et al.*, 2022). The regulation on ethylene-regulated *EXPB3*, auxin-regulated *XTH33* and *Cellulose synthase-like* (*CSL*)- *C12* was lost in *S. parvula*, whereas the binding of AREB/ABFs to *XTH18* and *CSLC5* was found. Transcriptomics analysis showed higher expression levels of auxin-regulated *XTH3, 18* and *33* in roots of *S. parvula* than Arabidopsis in response to ABA. Taken together, transcriptomic profiles and measurements on cortical cell length suggest that *S. parvula* can maintain cell expansion via both cell wall loosening and biosynthesis which finally may enable *S. parvula* to keep growing roots under severe salt stress.

## Acknowledgements

We are grateful for the help of Florian Galbier (ETH Zurich) and Dr. Michaela Fischer-Stettler (ETH Zurich) in radioactive lab work on ^14^C labelling, and Eva van Zelm and Jasper Lamers for sharing their scripts for root tracing. The visit of. H.L. to the laboratories of Dr. Diana Santelia and Prof. Samuel C. Zeeman was sponsored by an EMBO Short-Term Fellowship. H.L. is funded by the China Scholarship Council (CSC) through a Sino-Dutch Bilateral Exchange Scholarship (CF 13731). C. T. and Y.Z. are funded by the European Research Council (ERC) under the EU Horizon 2020 Research and Innovation programme (grant agreement 724321). C.P. is funded by Swiss National Science Foundation (SNF grant 310030_185241/1). Data produced in this work were partially generated in collaboration with the Genetic Diversity Centre, ETH Zurich.

## Author contributions

H.L. performed most experiments and their data analysis. K.D. contributed the RNA-seq data analysis. J.S. conducted root anatomical experiments and measurements of ions and performed the data analysis of these experiments. J.S. and N.W. performed halotropism assays. H.L., C.P., D.S. and S.C.Z. designed the ^14^C labeling and starch/sugar measurement experiments; H.L. and C.P. performed ^14^C labeling and starch/sugar measurements and data analysis. C.T. conceived the project, C.T and Y.Z. guided the research and C.T. H.L. and Y.Z. designed most experiments and wrote the manuscript. All authors commented on and approved the manuscript.

## Data Availability Statement

The data that support the findings of this study are available in the supplementary material of this article. The sequence data and supplementary tables that support the findings of this study will be openly available after acceptance.

## Supporting Information

**Table S1** RNA-seq results comparing the expression levels of genes between salt stress (175mM) and control conditions(0mM) in roots of *S. parvula* at 3 h, 24 h and 48 h under salt stress (separate XLSX).

**Table S2** Subgroups of differentially expressed genes (DEGs) that present in all timepoints, in 2 timepoints or specifically in one timepoint in roots of *S. parvula* under salt stress (separate XLSX).

**Table S3** Gene ontology (GO) enrichment analysis on DEGs that are down- or up-regulated at 3 h, 24 h or 48 h in roots of *S. parvula* under salt stress (separate XLSX).

**Table S4** GO enrichment analysis on subgroups of DEGs shown in Table S2 (separate XLSX).

**Table S5** Gene list of clusters from comparative analysis via k-means clustering (separate XLSX).

**Table S6** GO enrichment analysis on genes of *S. parvula* clustered via k-means clustering shown in Table S5 (separate XLSX).

**Figure S1.**
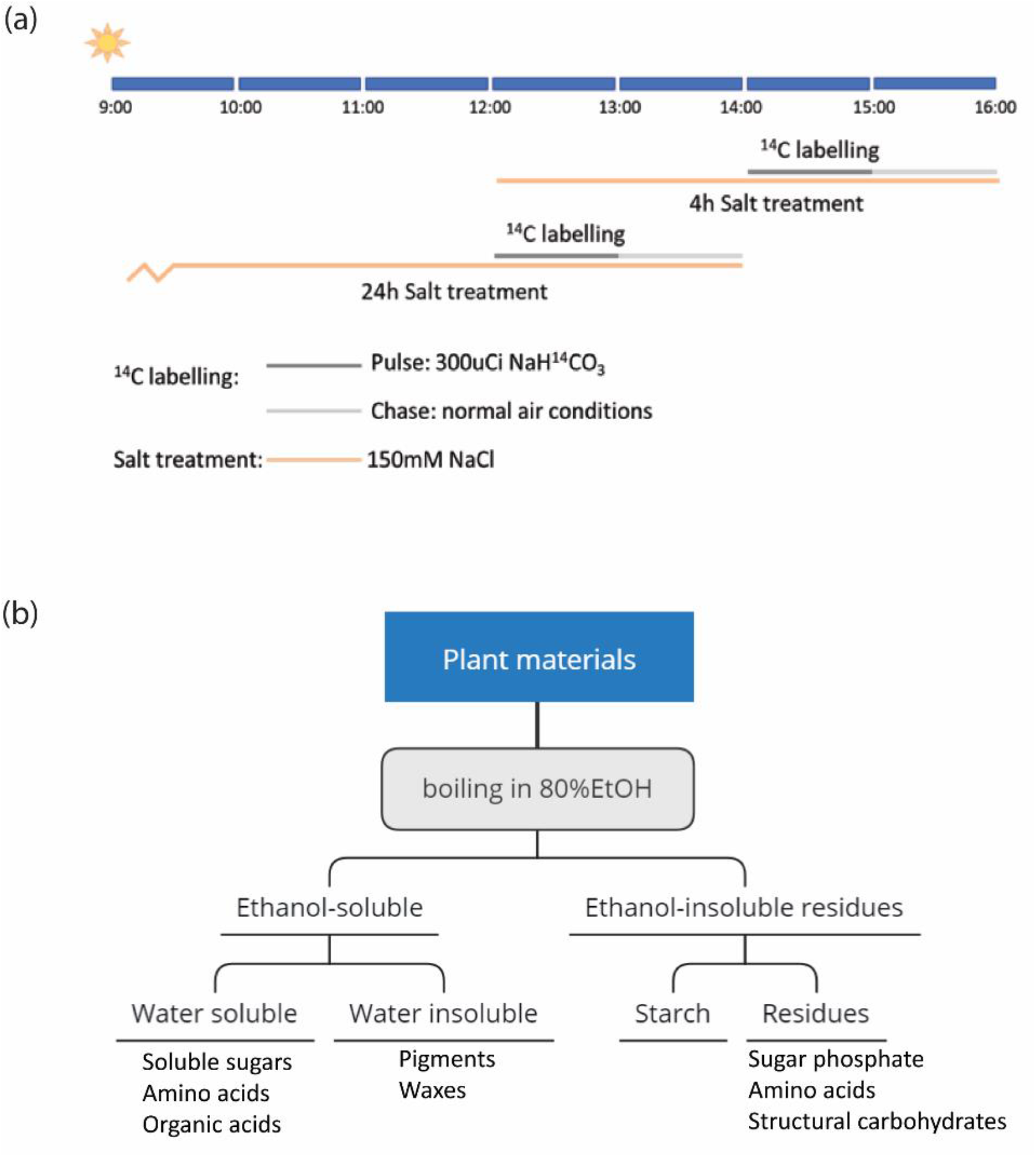
Scheme of ^14^CO_2_ Pulse-Chase Labeling setups and sample processing. (a) Scheme of ^14^CO_2_ Pulse-Chase Labeling setups. 10-day-old seedlings of *S. parvula* and Arabidopsis grown on ½ MS medium were transferred to ½ MS medium supplemented with 0 mM or 150 mM NaCl for 4 h or 24 h. plants were pulsed with 300 μCi ^14^CO_2_ for 60min in a sealed Plexiglas chamber at 2 h before the end of the salt treatments and then chased in normal air for 60min. (b) Sample processing. Labelled shoots and roots were harvested and pooled separately. Samples were boiled in 3 mL 80% ethanol for 10min and homogenized. Supernatant and pellet were separated by centrifuging at 2400g for 12min. Pellet was washed sequentially with 1 mL of 50% ethanol, 20% ethanol, H_2_O and 80% ethanol. For each replicate, all the supernatants from the washing steps were pooled together and resuspend in 2 mL of water after being dried in vacuum, yielding the water-soluble compounds. 1 mL of 100% ethanol was subsequently used in the same tube to obtain the ethanol soluble compounds which include pigments, waxes and lipids. Neutral sugars in the water-soluble fraction were eluted through home-made 1.5 mL columns of cation exchanger Dowex 50 W and anion exchanger Dowex 1 (Sigma-Aldrich). The pellets from the root samples, containing the insoluble compounds, were air-dried and resuspended in 1 mL of Soluene™-350 (PerkinElmer) for 24 hours, before measuring. Pellets from the shoot samples were resuspended in 1 mL of water and a 100 ul aliquot was further digested by amyloglucosidase (Roche) and α-amylase (Roche) in a 9:1 ratio to measure the starch content (hydroyzed to water-soluble glucose) and the insoluble compounds (resuspended in 1 mL of Soluene™-350 after 15 min centrifugation at 5000g). The radioactivity of ^14^C incorporated in each fraction in disitegrations per minute (dpm), was measured in a total volume of 5 mL with a LS1801 scintillation counter (Beckmann) as described in (Kolling *et al.*, 2013), by adding 4 mL of Ultima Gold™ (PerkinElmer) for water soluble fractions or Ultima Gold™ LLT (PerkinElmer) for ethanol/ Soluene™-350 fractions, to 1 mL of sample. The amount of radioactivity present in each fraction was expressed as the ratio of dpm in each fraction over the sum of dpm in all fractions for a given sample.

**Figure S2.**
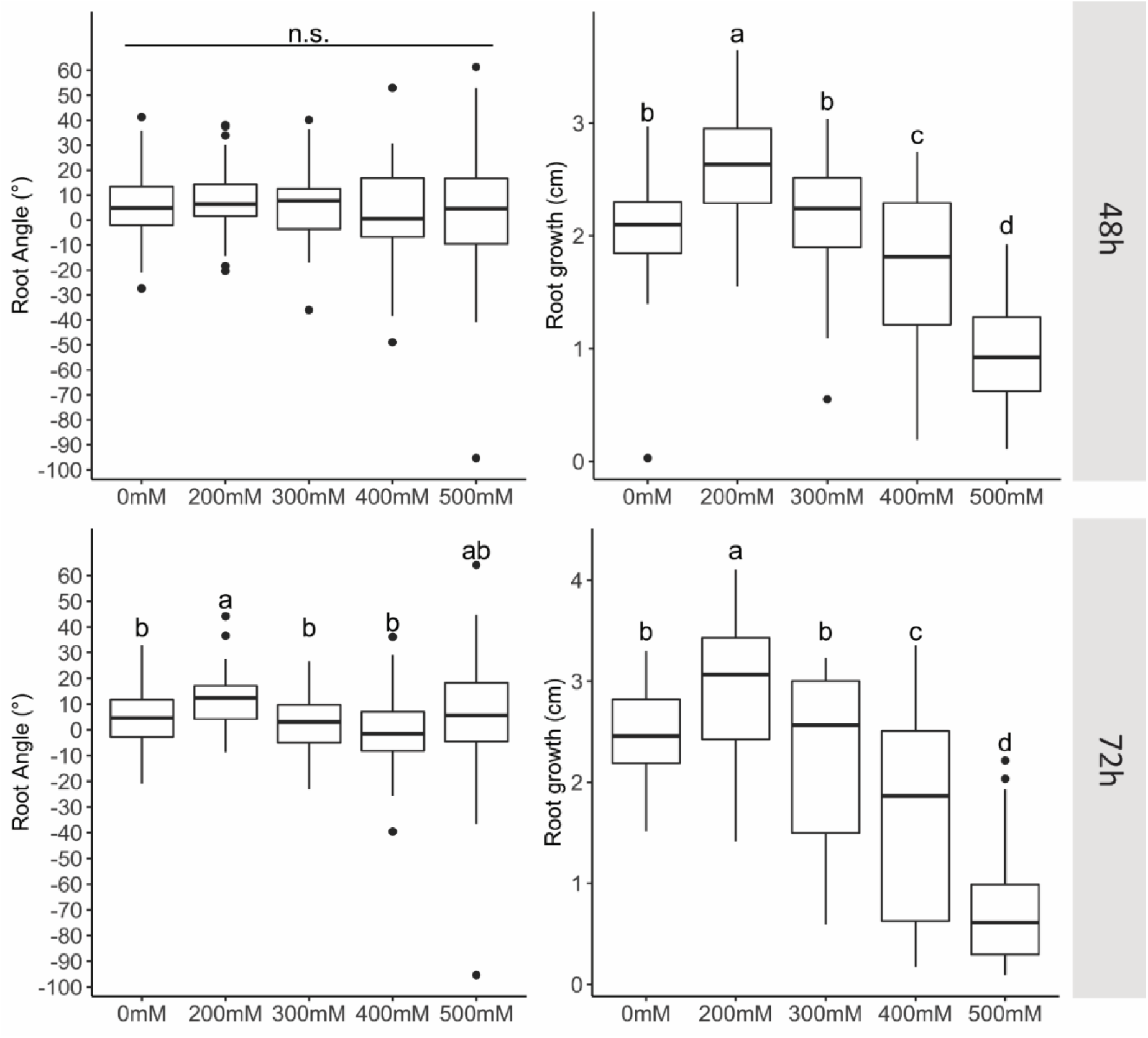
*S. parvula* roots do not exhibit a halotropic response. 4-day-old *S. parvula* seedlings germinated and grown on ½ MS agar plates supplemented with 0.5% sucrose were transferred to replacement medium 48 h or 72 h (n=63-71). Replacement medium (lower diagonal half) consisted of ½ MS medium supplemented with 0.5% sucrose and with 0 mM, 200 mM, 300 mM, 400 mM or 500 mM NaCl. Significant differences (p<0.05) indicated by letters were determined by two-way ANOVA followed by Turkey’s post-hoc test and “n.s.” denotes no significant differences.

**Figure S3.**
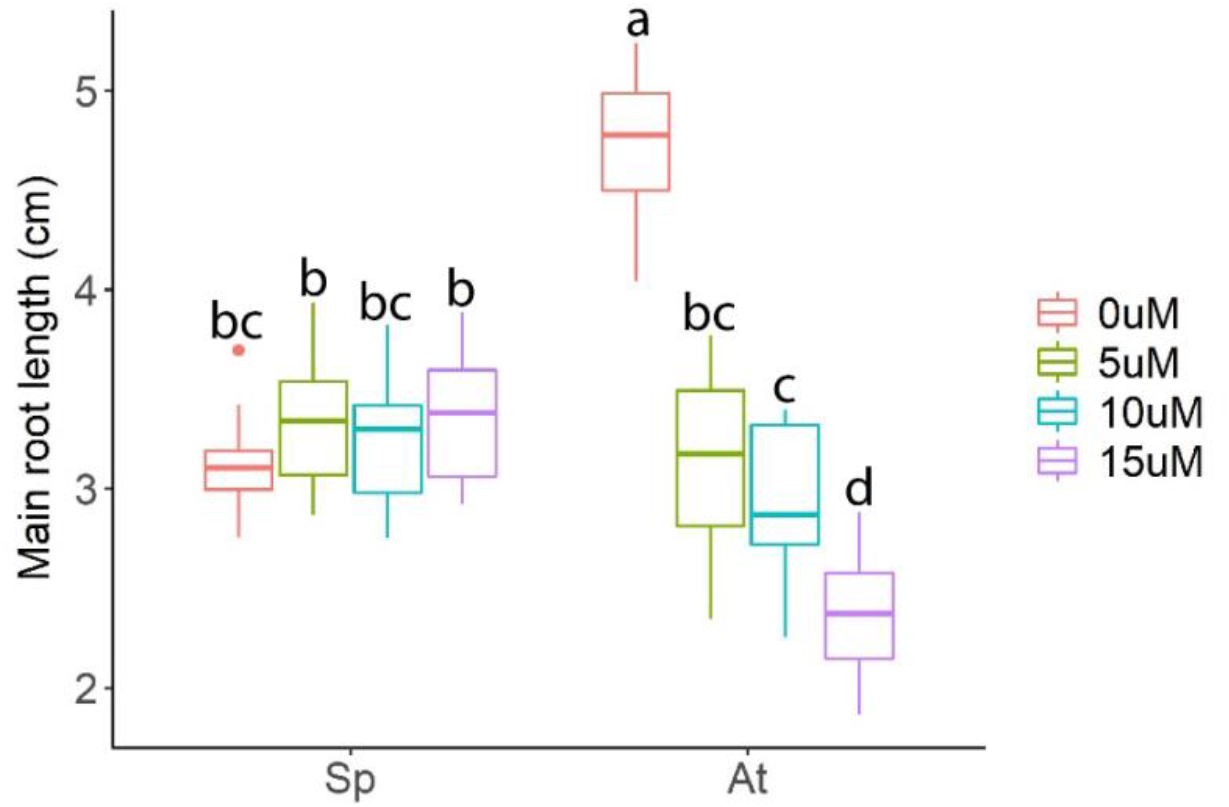
Primary root growth of *S. parvula* is less inhibited by ABA compared to Arabidopsis. 4-day-old seedlings of *S. parvula* and Arabidopsis were transferred to ½ MS medium supplemented with 0μM, 5μM, 10μM, or 15μM ABA for 6 days (n=10-20). Significant differences (p<0.05) indicated by letters were determined by two-way ANOVA followed by Turkey’s post-hoc test.

**Figure S4.**
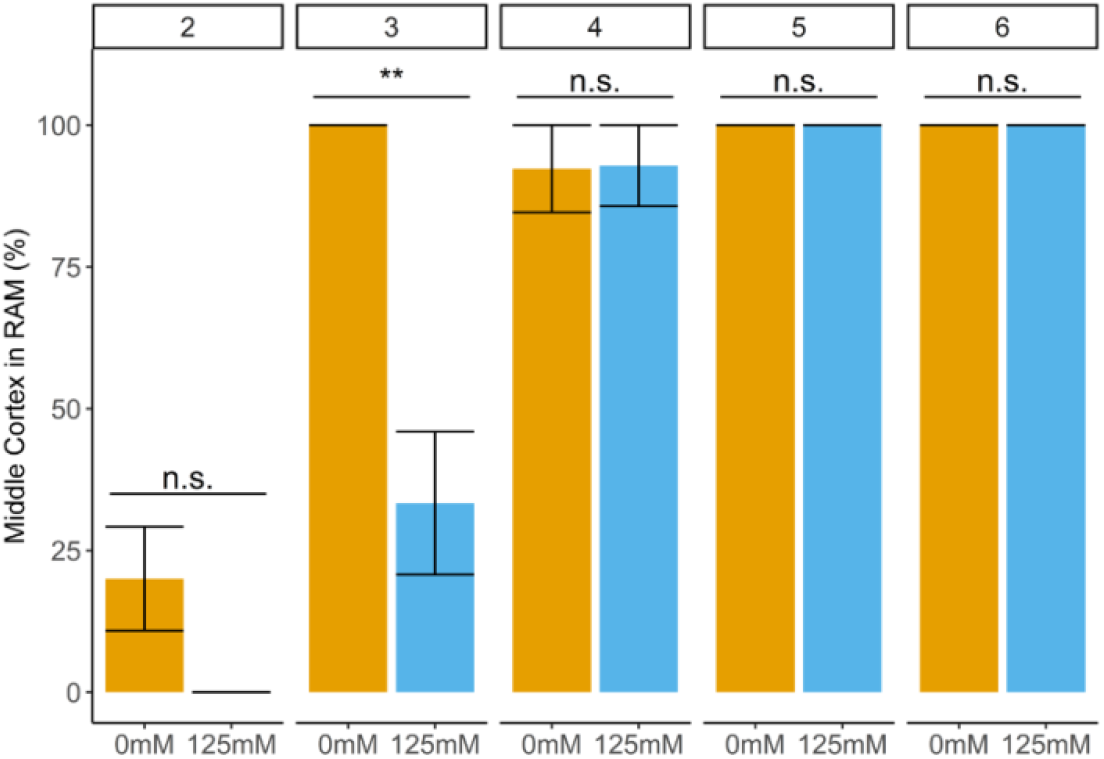
Salt delays middle cortex formation in Arabidopsis. 4-day-old seedlings of Arabidopsis germinated and grown on ½ MS agar plates were transferred to ½ MS agar plates supplemented with 0 mM or 125 mM NaCl for 1 to 6 days (upper bar). The presence of middle cortex was checked in root apical meristem of Arabidopsis seedlings. 0% denotes absence of MC in all samples and 100% denotes presence of MC in all samples (n=13-25). Statistical comparisons by Chi-Squared Test (p <0.05) where “n.s.” denotes no significant differences and “ ** ” denotes significant difference with p-value = 0.001559.

**Figure S5.**
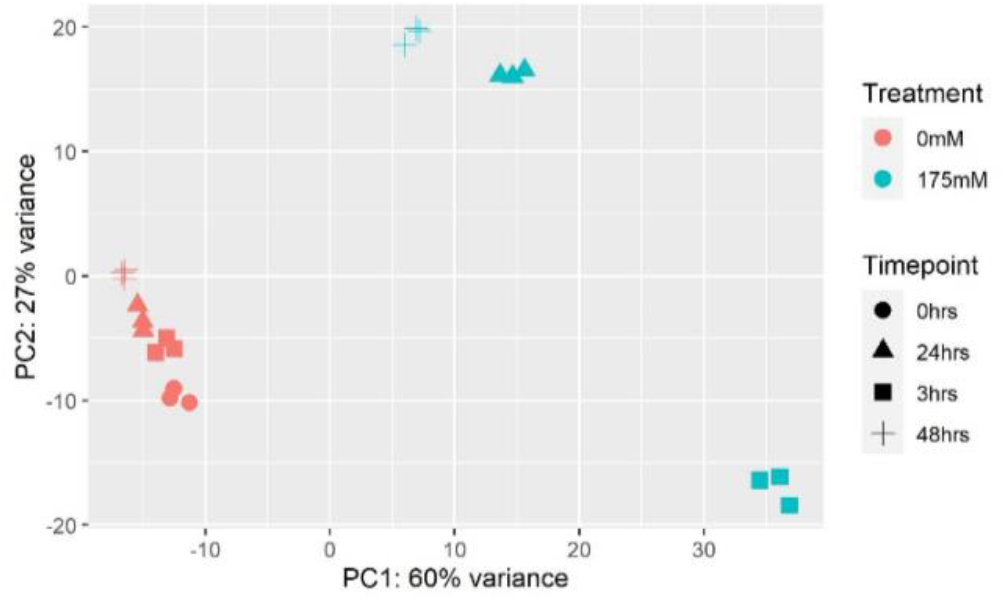
Principal component analysis of the different samples revealed a temporal response to salt. Normalized counts of all transcripts under 0 mM and 175 mM NaCl were analyzed. The first two components are presented. Colors indicate the different treatments and shapes indicate the timepoints after treatments.

**Figure S6.**
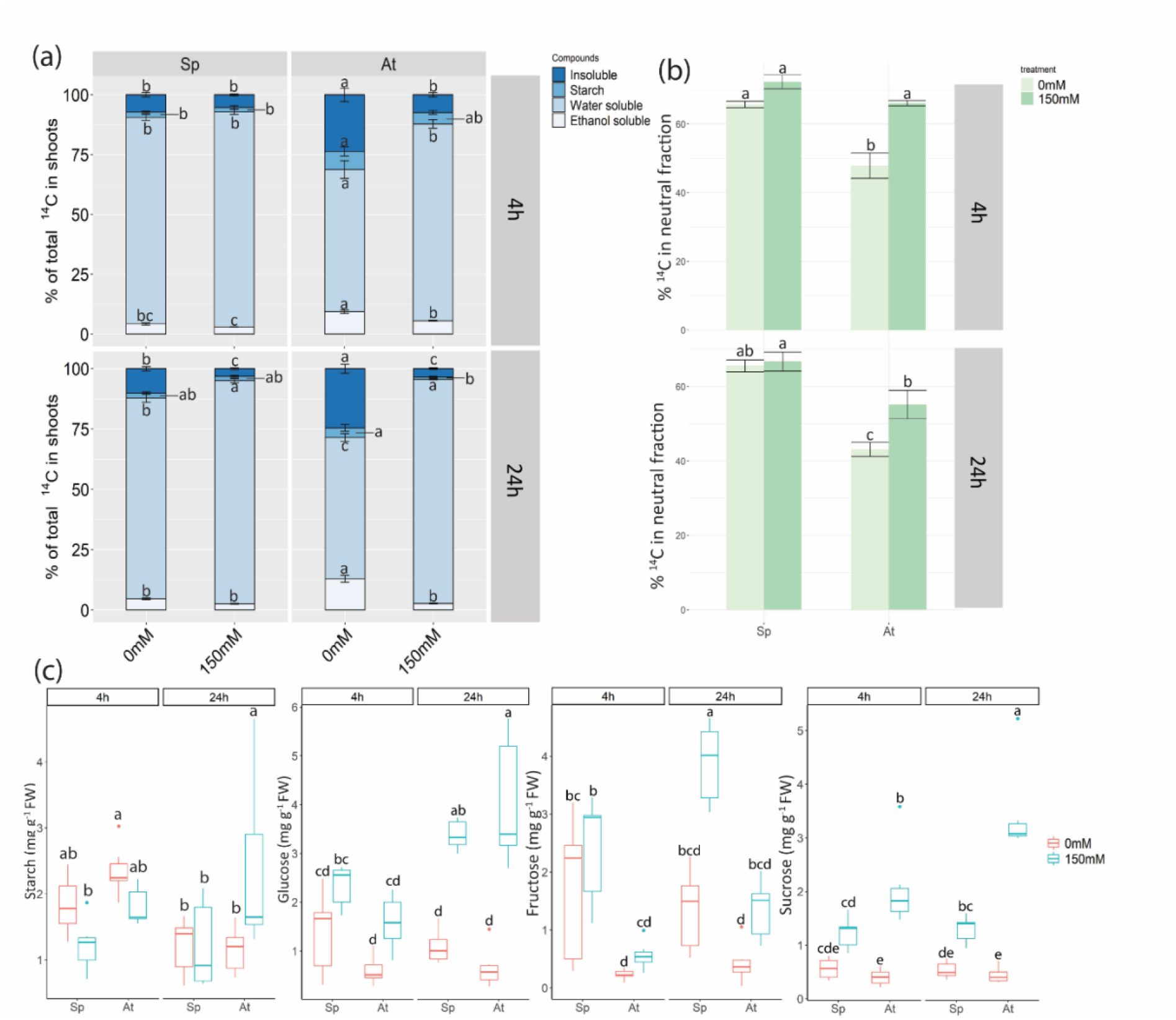
Impact of salt stress on carbon allocation in shoots of *S. parvula* compared with Arabidopsis. 10-day-old seedlings of *S. parvula* and Arabidopsis grown on ½ MS medium were transferred to ½ MS medium supplemented with 0 mM or 150 mM NaCl for 4 h or 24 h. ^14^CO2 pulse-chase experiment was performed on plants exposed to salt stress for 4 h and 24 h (n= 5-6 pools of 10-15 shoots or roots). (a) The incorporation of ^14^C into different compounds in shoots after 4 h and 24 h under salt stress. (b) The amount of incorporated ^14^C into the soluble sugars in shoots after 4 h and 24 h exposure to salinity were given as percentages of incorporated ^14^C in water-soluble compounds. Error bars represent standard error. (c) measurements of starch, soluble sugars (glucose, fructose and sucrose) content in shoots of *S. parvula* and Arabidopsis after 4 h or 24 h under salt stress. FW, fresh weight. Significant differences (p<0.05) indicated by letters were determined by two-way ANOVA followed by Turkey’s post-hoc test.

**Figure S7.**
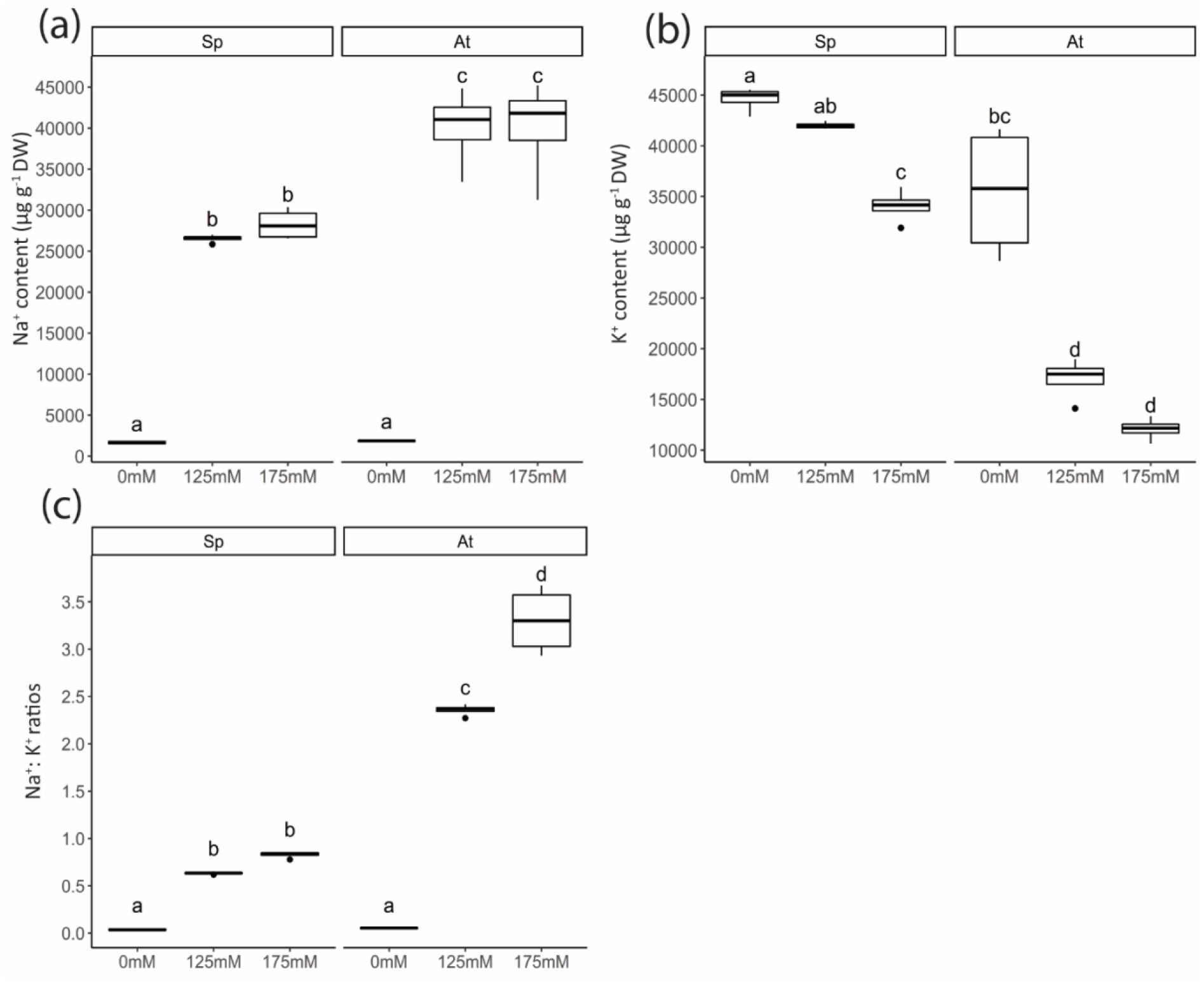
Better maintenance of Na^+^:K^+^ ratios in *S. parvula* seedlings under salt stress. Less sodium accumulation (a) and higher potassium levels (b) in response to salt results in lower Na^+^:K^+^ ratios (c) in *S. parvula* than in Arabidopsis. 4-day-old seedlings of Arabidopsis and *S. parvula* germinated and grown on ½ MS agar plates were transferred to ½ MS agar plates supplemented with 0 mM, 125 mM or 175 mM NaCl for 2 days (n= 4 pools of approx. 70 plants). Error bars represent standard error. Significant differences (p<0.05) indicated by letters were determined by two-way ANOVA followed by Turkey’s post-hoc test.

**Figure S8.**
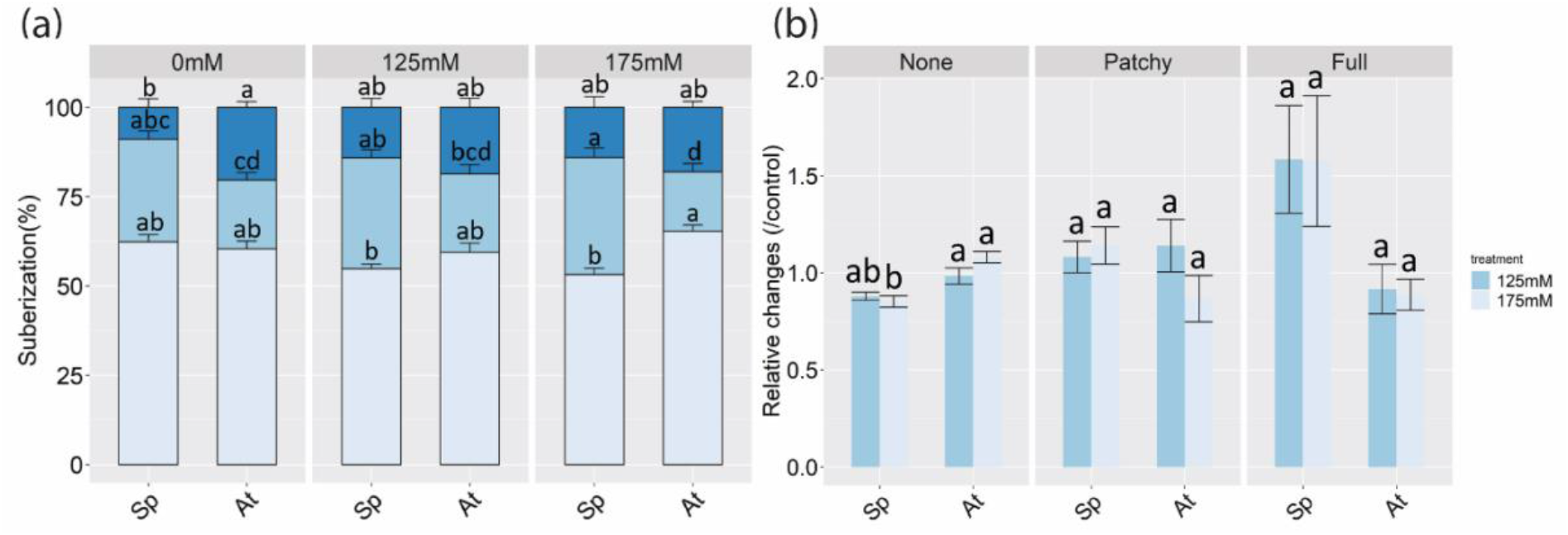
Suberization did not show significant changes in roots of *S. parvula* and Arabidopsis after 6 h exposure to salt. 4-day-old seedlings of *S. parvula* and Arabidopsis were transferred to ½ MS sucrose medium supplemented with 0 mM, 125 mM or 175 mM NaCl for 6 h and suberin was stained by FY 088. Suberization was separated into no-suberized zone (None), patchy (Patchy) and continuous suberization (Full). (a) Quantification of suberin deposition along the root axis and was indicated with three different zones (n=23-27). Data are presented as percentages of root length. Error bars represent standard error. Significant differences (p<0.05) indicated by letters were determined by two-way ANOVA followed by Turkey’s post-hoc test. (c) Quantification of relative changes of 3 suberization zones under salt stress. Data are calculated by the ratio between salt treatment and control condition. Error bars represent standard error. Significant differences (p<0.05) indicated by letters were determined by two-way ANOVA followed by Turkey’s post-hoc test.

**Figure S9.**
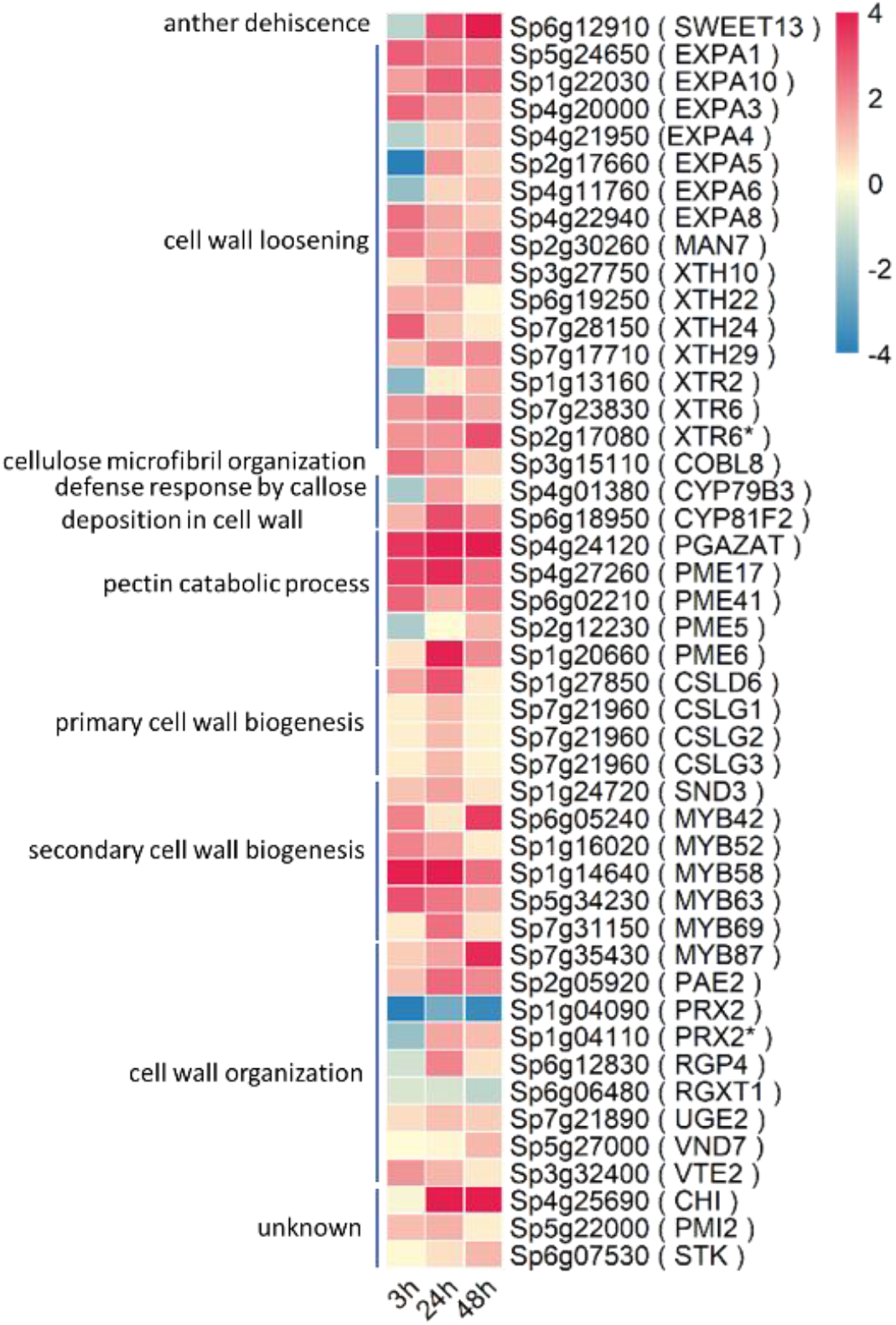
Heatmap presenting expression profiles of DEGs enriched in GO term ‘Cell wall organization or biogenesis’ at 24 h and 48 h in roots of *S. parvula* under salt stress. Gene expressions were extracted from transcriptomic data analysis (p< 0.05 Log2 Fold changes >1). Gene symbols based on the annotations of Arabidopsis were shown in brackets (rows). Colors indicate the Log2 Fold changes between −4 and 4.

**Figure S10.**
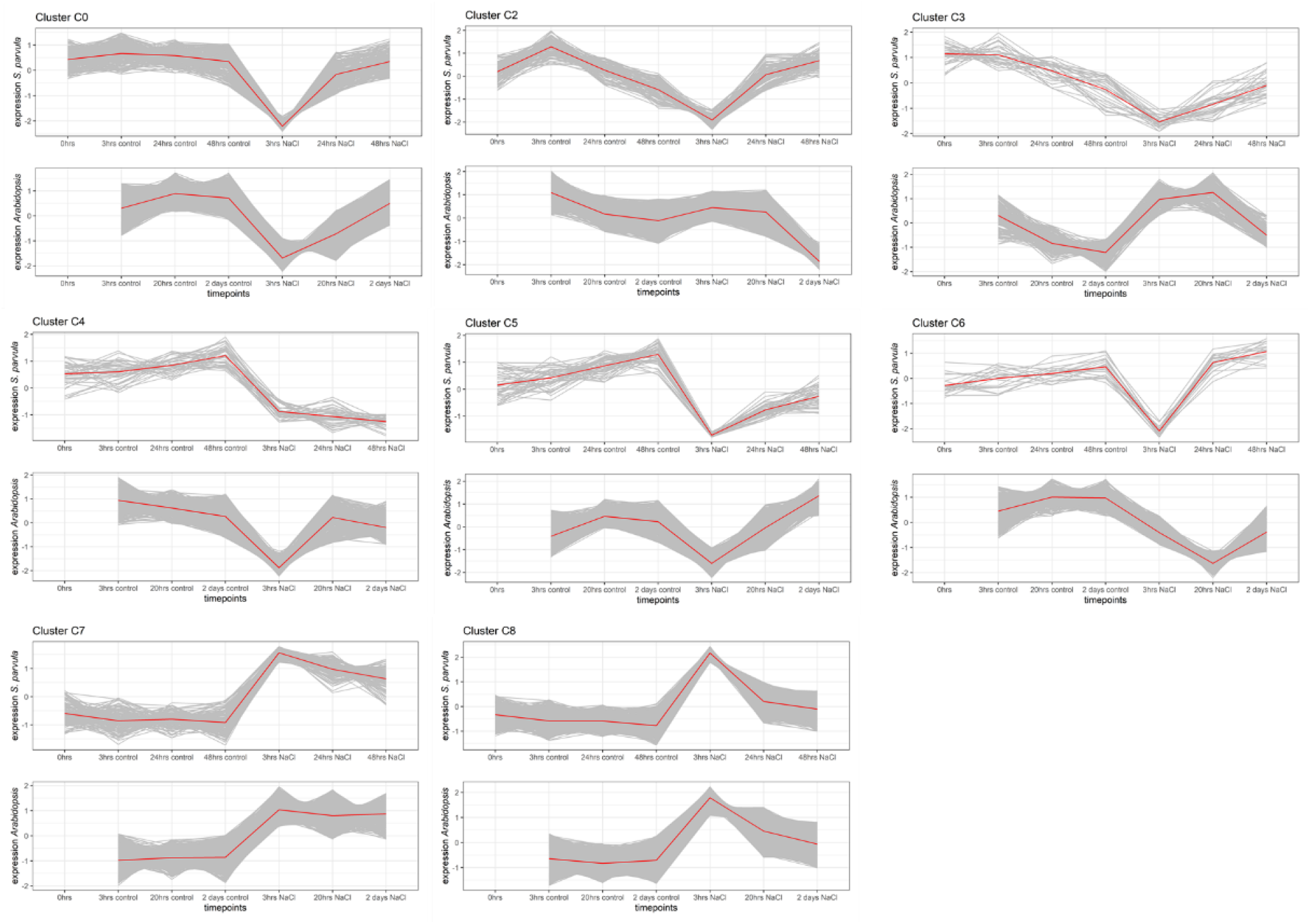
Comparative analysis on transcriptomes of roots of both Arabidopsis and *S. parvula* were performed using k-means clusters. FPKM values from the dataset of *S.parvula* and normalized microarray expression data of timepoints (3 h, 20h and 2 days) of Arabidopsis were used (Geng *et al.*, 2013).

**Figure S11.**
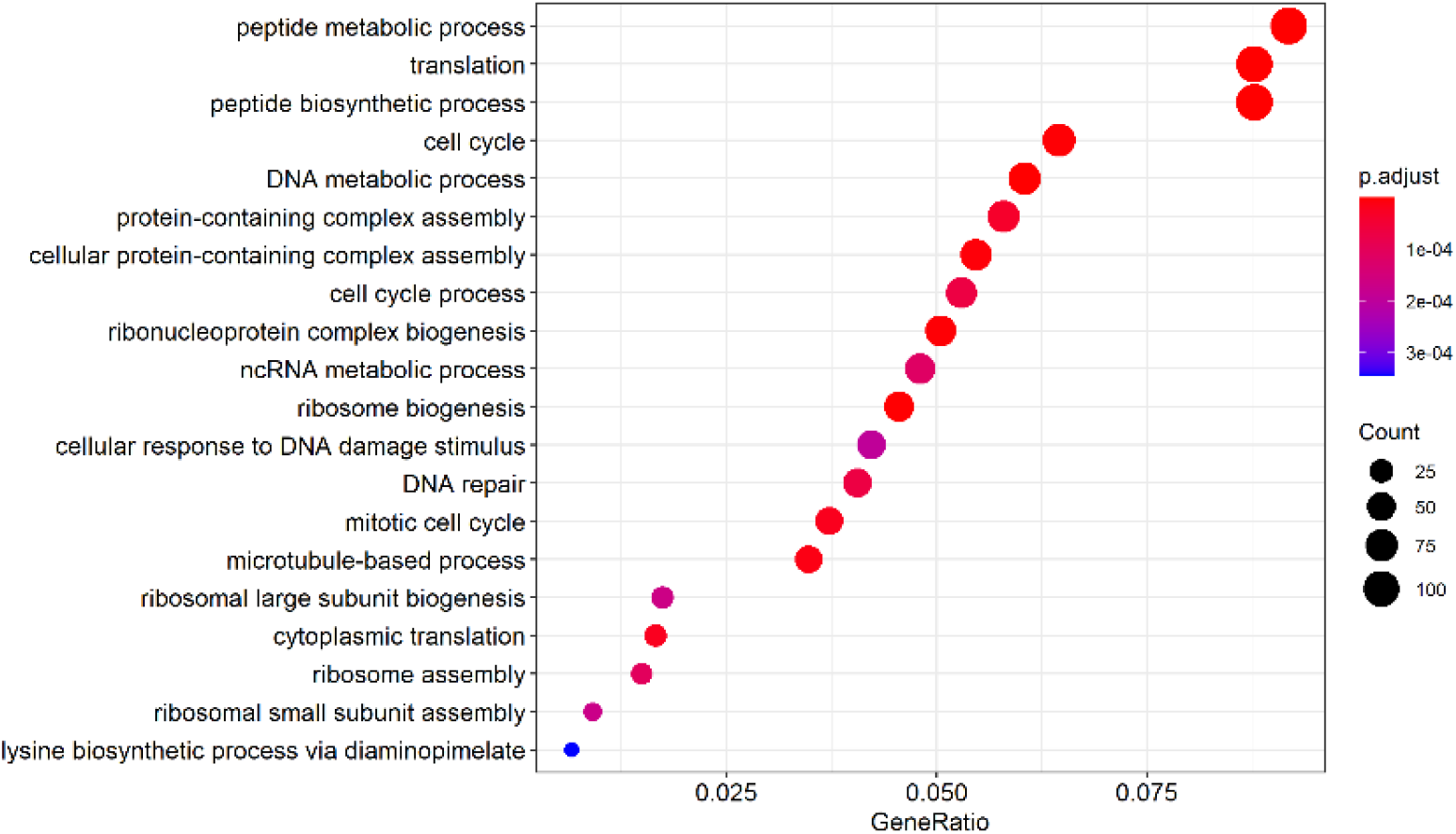
Gene ontology (GO) enrichment analysis of genes from cluster 1 based on biological process (BP).

**Methods S1** Ion measurements

Four-day-old seedlings of Arabidopsis and *S. parvula* were transferred to agar plates supplemented with 0mM, 125mM or 175mM NaCl for 2 days. Four pools of 70 seedlings were harvested and dried at 80°C for one day. Ion content was measured using ICP-MS at the Ionomics Facility, University of Nottingham, United Kingdom(Danku *et al.*, 2013).

**Methods S2** Primary root growth measurements

Seeds of *S. parvula* were germinated on ½ MS agar plates and transferred after 4 days to new plates supplemented with 0μM, 5μM, 10μM or 15μM ABA for 6 days. Plates with 10-day-old seedlings were scanned with an Epson perfection V800 scanner at 400 dpi. Scanned images were pre-traced by an automated script based on root edge detection (https://github.com/jasperlamers/RootFinder/tree/main). Manual correction was performed with an ImageJ plugin Smartroot (Lobet *et al.*, 2011).

